# FrustratometeR: an R-package to compute local frustration in protein structures, point mutants and MD simulations

**DOI:** 10.1101/2020.11.26.400432

**Authors:** Atilio O. Rausch, Maria I. Freiberger, Cesar O. Leonetti, Diego M. Luna, Leandro G. Radusky, Peter G. Wolynes, Diego U. Ferreiro, R. Gonzalo Parra

## Abstract

Once folded, natural protein molecules have few energetic conflicts within their polypeptide chains. Many protein structures do however contain regions where energetic conflicts remain after folding, i. e. they have highly frustrated regions. These regions, kept in place over evolutionary and physiological timescales, are related to several functional aspects of natural proteins such as protein-protein interactions, small ligand recognition, catalytic sites and allostery. Here we present FrustratometeR, an R package that easily computes local energetic frustration on a personal computer or a cluster. This package facilitates large scale analysis of local frustration, point mutants and molecular dynamics (MD) trajectories, allowing straightforward integration of local frustration analysis into pipelines for protein structural analysis.

**Availability and implementation:** https://github.com/proteinphysiologylab/frustratometeR

## Introduction

Proteins are evolved biological molecules that adopt a defined set of structures constituting their ‘native’ state. Built as linear polymers, proteins find their native state easily since evolution has minimized the internal energetic conflicts within their polypeptide chain, following the “principle of minimal frustration” (1). Proteins are not only biologically optimized to fold or to be stable but also to ‘function’ (2, 3) and therefore it’s not surprising to find that about 10-15% of the internal interactions in proteins are in energetic conflict within their local structure (4). These conflicts, kept in place over evolutionary and physiological time scales, allow proteins to explore different conformations within their native ensemble and thus enable the emergence of ‘function’. Over the last years, the concept of local frustration has given insights into a diverse set of functional phenomena: protein-protein interactions, ligand recognition, allosteric sites (5), enzymatic active sites and co-factors binding (6), evolutionary patterns in protein families (7), polymorphisms in the Class I Major Histocompatibility Complex (MHC) (8), disease associated mutations (9), protein dynamics (10), disorder-to-order transitions in protein complexes (11, 12) as well as fuzzy interactions (13) and chaperones-clients recognition (14). Frustration and its role in functional dynamics has been recently reviewed (15). Up to now, facile location and quantification of energetic frustration has been made possible via the Frustratometer web server (16, 17). Unfortunately high-throughput analysis using the server is not feasible as the flexibility of the algorithm was reduced in order to maximise usability by non computational scientists. Here we present FrustratometeR, an R package that retains all the capabilities present in the web server but that also includes brand new modules to evaluate how frustration varies upon point mutations as well as to analyse how frustration varies as a function of time during molecular dynamics simulations. FrustratometeR facilitates the analysis both at small and high-throughput scales and thus allows one to integrate local frustration analysis into other protein structural bioinformatics pipelines.

## Methods

Full methods descriptions are available in the supplementary material. Fig1 summarises FrustratometeR functionalities and minimum code to generate plots there in (Fig1E). Here we describe the main functions in the package. The three functions can be used in combination with any of the frustration indexes (i.e. *configurational, mutational or singleresidue*). If not specified the *configurational* mode is used as default.

**Fig 1.**
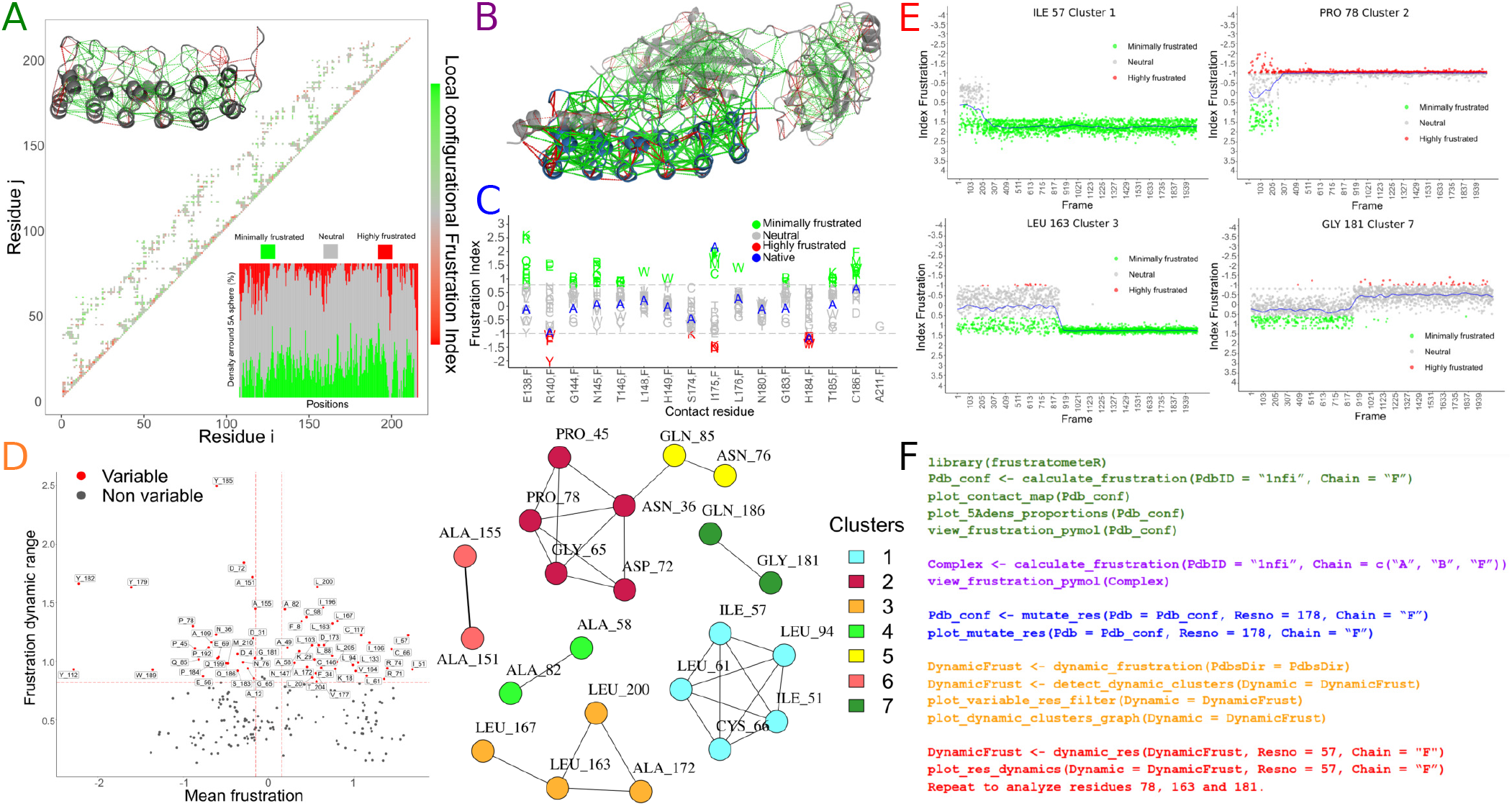
Frustration in I*κ* B*α* (PDB ID 1nfi,F). (A) Contact map, 5Adens plot and pymol representation. (B) Frustration of I*κ*B*a* in complex with Nf*κ*B (chains A,B,F). (C) Frustration changes when mutating a specific residue to all canonical amino acids alternatives. X axis: all residues that interact with the residue of interest for all mutants are displayed. Y axis: frustration values are shown and coloured based on their frustration classification. Native variant appears in blue. Each mutant is represented by its 1-letter amino acid code to identify to which variant it corresponds. (D) Dynamic frustration modules. Left: Residues that vary their frustration across frames in molecular dynamic simulations are identified based on their frustration average and dynamic range values. Right: Variable residues are connected to each other in a correlation network that is then clustered to find modules with similar dynamic behaviour (details in supplementary material). (E) Frustration values as a function of time (simulation frames) for representative residues in Clusters 1-4. (F) Minimum code to generate the previous panels

### Calculate local frustration

A single function, *calculate_frustration()*, the PDB structure or PDBID and the desired frustration index specification are enough to calculate the local frustration either for single (Fig1A) or multiple chain structures (Fig1B). Optionally an electrostatic term can be activated in the energy function by giving a value to the *Electrostatics_K* parameter that is set by default to *NULL* (see supplementary material or (17) for details).

### Frustration upon mutations

The *mutate_res()* function allows users to analyse how point mutations affect local frustration (Fig1C). For every possible alternative amino acid, this function generates a structural variant. Two modes are implemented to generate the mutants: *threading* (it does not modify the backbone coordinates) and *modeller* (performs some energetic optimization by generating an homology model with Modeller (18)). Subsequently, frustration for each mutant is calculated as with *calculate_frustration()*.

### Frustration along MD simulations

The *dynamic_frustration()* function analyses frustration along a molecular dynamics trajectory (Fig1D). The *detect_dynamic_clusters()* function can group protein residues based on their temporal “singleresidue” frustration profiles and find dynamic modules constituted by highly correlated residues (Fig1D). Individual residues temporal dynamics can be visualized as well (Fig1E).

## Discussion

As an example, we have applied FrustratometeR to I*κ*B*α*, an inhibitor of Nf*κ*B (PDBID 1nfi). Fig1A shows a composite figure of typical Frustratometer web-server-like visualisations (17) for I*κ*B*α* (chain F). Fig1B shows frustration results for I*κ*B*α* in complex with Nf*κ*B (chains A, B, F). Bear in mind that frustration results differ when chains from quaternary complexes are analysed in isolation or in complex due to the existence of compensating interactions between interacting partners. Far from being a disadvantage, differences between monomers and quaternary complexes frustration results can be (and have been) exploited to analyse interaction mechanisms (7). FrustratometeR introduces a new functionality to evaluate the change in frustration for amino acid variants which can be used as a guide to tune specific residue-residue interactions in the structure. Fig1C shows the change in the configurational frustration index for residue ALA178 when it is mutated to all the other amino acids in *threading* mode. FrustratometeR also includes a module to analyse molecular dynamics simulations and to identify residues that have similar dynamics. We extracted frames from an I*κ*B*α* coarse-grained folding simulation (see supplementary data for details). First, *m* variable residues are selected based on their average and the dynamic range of “singleresidue” frustration values across frames. A low dimensional representation of a matrix of *m* residues and *n* frames is obtained using Principal Components Analysis (PCA) and residues pairwise correlations are calculated to create a graph that is clustered to define dynamic modules (Fig1D). Here we have analysed the trajectory by detecting clusters with adjusted parameters (CorrType = “spearman”, FiltMean = 0.15, MinFrstRange = 0.7, MinCorr = 0.95; see supplementary material for more detailed explanations). Although a detailed analysis of the folding mechanism of the I*κ*B*α*/Nf*κ*B is beyond the scope of this article, even this simple analysis can give very useful insights about the system. After filtering residues and building a graph with the dynamic residues, seven clusters are left (Fig1D). I*κ*B*α* folding is coupled to Nf*κ*B recognition, it involves disorder/order transitions and its energy landscape is quite complex (see “detect dynamic clusters” section in supplemental material for an energy landscape representation). In a very condensed fashion, we can say that there are two major transitions: 1) from the unfolded state to an inter-mediate state when I*κ*B*α* N-terminal repetitive region starts to interact with Nf*κ*B and 2) from the intermediate state to an expanded/folded state when the C-terminal repeats from I*κ*B*α* get structured by forming further stable interactions with Nf*κ*B (19). Clusters 1 and 2 involve residues that have a frustration change during the first folding transition (around frame 200). Cluster 1 residues change their state from being neutral or highly frustrated to become minimally frustrated and are mostly located within helices in repeats 1 and Cluster 2 residues change their state from being minimally frustrated to become highly frustrated. These residues are involved in the interactions with Nf*κ*B and become frustrated because the structure is forced to adopt its quaternary structure conformation without Nf*κ*B being present to energetically compensate them. Clusters 3 and 7 behave much like clusters 1 and 2 but their frustration changes occur during the second folding transition (around frame 850). A more detailed analysis is made in the *detect dynamic clusters* section in supplemental material.

## Conclusion

We present a user-friendly R package to calculate energetic local frustration in protein structures. The package includes new features to assess the effect of mutations in local frustration as well as to analyse frustration along molecular dynamics trajectories. Its simple interface together with the newly implemented functionalities will facilitate frustration analysis at larger scales and can be used to include FrustratometeR as part of different pipelines for protein structural analysis.

## Supporting information

Supplementary Material - Vignette.

