## Supplementary Material - Vignette. for "FrustratometeR: an R-package to compute local frustration in protein structures, point mutants and MD simulations"

[1]Facultad de Ingeniería, Universidad Nacional de Entre Ríos, Argentina.
[2]Laboratorio de Fisiología de Proteínas, Departamento de Química Biológica -IQUIBICEN/CONICET, Facultad de Ciencias Exactas y Naturales, Universidad de Buenos Aires, Argentina.
[3]Center for Genomic Regulation, Barcelona Institute for Science and Technology, Barcelona, Spain.
[4]Center for Theoretical Biological Physics and Department of Chemistry, Rice University, Houston, TX, 77005, USA.
[5]European Molecular Biology, Laboratory, Heidelberg, Germany.
[*]Jointly 1st authors

### 1. Description

### FrustratometeR: An R package to calculate energetic local frustration in proteins

Energetic local frustration has been extensively linked to multiple functional aspects of proteins. The protein frustratometer has been present as a web server service since 2012. Here we present, frustratometeR, a standalone R package that extends the set of analysis present at the web server together with brand new functionalities to help elucidate the role of local frustration in proteins function and dynamics.

**Description:**

Given a PDB file and a frustration index type, several visualisations and pymol scripts are produced. Additionally the frustratometeR package can compute the frustration index distribution for all alternative amino acids for a given residue to evaluate the impact of point mutations in the structure. A module to analyze frustration change in Molecular Dynamics simulations is implemented.

This standalone version of the frustratometer has long been expected by many users. The frustratometeR not only allows to perform frustration calculations locally but also extends its functionalities to study the impact of residue mutations and the mechanistic role of frustration during protein dynamics.

### 2. Installation

#### Installation

`invisible(lapply(c("usethis", "devtools"), library, character.only = TRUE))`

`devtools::install_github("proteinphysiologylab/frustratometeR")`

#### Dependencies

##### R packages (version R >= 3.6.3 required)

`install.packages(c("usethis", "devtools"))`

##### Python 3

- numpy `python3 -m pip install numpy`
- biopython `python3 -m pip install biopython`
- leiden(optional) `python3 -m pip install leidenalg`

##### Others

- pymol `sudo apt install pymol`
- magick `sudo apt-get install -y libmagick++-dev`
- perl `sudo apt-get install perl`
- modeller (you need to install a licenced version of modeller in order to use some of the frustratometeR functionalities, https://salilab.org/modeller/) modeller needs to be available via python3. To check if this is working execute python3 and try to do "form modeller import * " after installing modeller. We suggest to install modeller using conda environments.

#### Minimum code to calculate frustration in a protein

`library(frustratometeR)`

`Pdb_conf <- calculate_frustration(PdbID = "1n0r",Chain = "A", ResultsDir = "/Home/Desktop/")`

`view_frustration_pymol(Pdb_conf)`

##### You can find an example of how to use the package at:

https://github.com/proteinphysiologylab/frustratometeR/tree/master/Examples

##### You can also find useful examples in our wiki!!:

https://github.com/proteinphysiologylab/frustratometeR/wiki

##### You can find the data used in the Wiki examples at:

https://github.com/proteinphysiologylab/frustratometer-data.git

### 3. Frustration calculation

#### Introduction to the calculation and analysis of local energy frustration in proteins.

The following diagram shows that the `calculate_frustration()` function calculates the local energy frustration from a *PdbFile* or *PdbId* of a protein structure, creating a Pdb Frustration Object that is used to generate visualisations, obtain frustration data and other processes how to analyze punctual mutations.

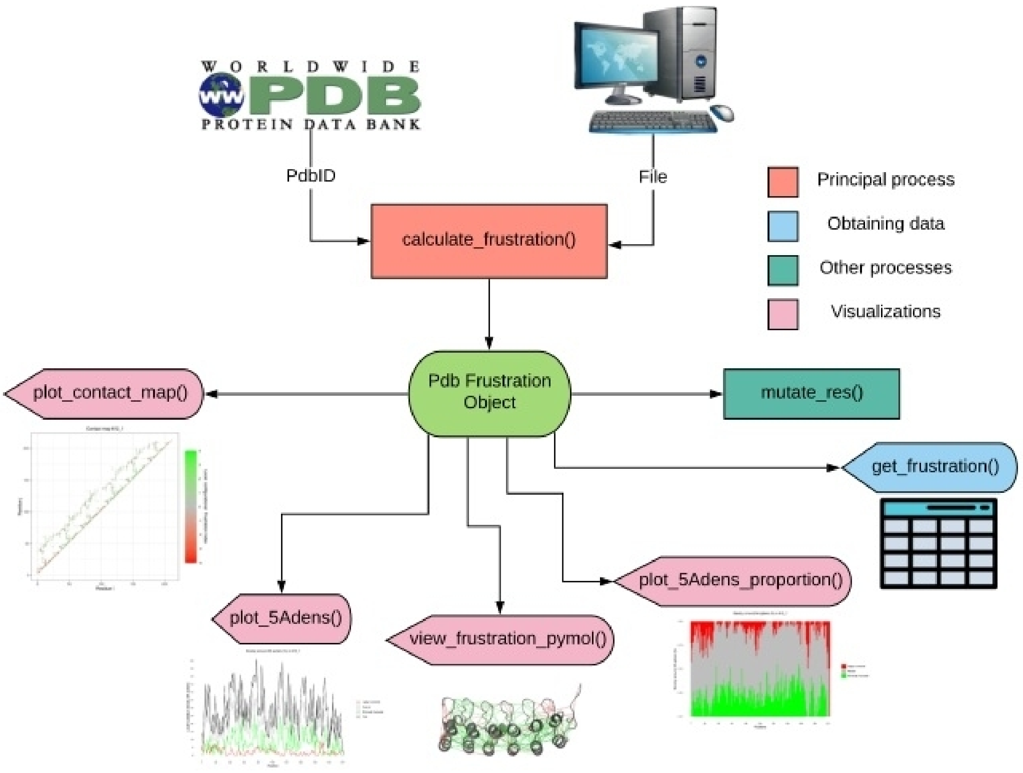


#### Example of analysis and explanation of use!

##### Load the frustratometeR library

```
library(frustratometeR)
```

Calculate the frustration of a structure to be downloaded from the *PDB*. If you have the structure, you can indicate its path with the *PdbFile* parameter. The calculation mode (**configurational**, **mutational** or **singleresidue**) can be indicated. The results are stored in *ResultsDir*. In addition, graphics may or may not be generated and stored at the output folder with the *Graphic* parameter. On the other hand, if you want to analyse only one chain (in case of multimeric structures) or multiple chains can be indicated with the *Chain* parameter.

```
Pdb_conf <- calculate_frustration(PdbID = "1nfi", Chain = "F", Mode = "configurational")
```

The energetic local frustration results can be obtained for processing as a Data Frame with the `get_frustration()` function. You can indicate a specific residue (*Resno*) or chain (*Chain*):

```
Frustration_data <- get_frustration(Pdb = Pdb_conf, Resno = 178, Chain = "F")
```

**The calculate_frustration() function can receive further parameters:**

_Electrostatics_K_: This is a numeric value that is set to NULL by default. If a numeric value is used a new term in the hamiltonian, that will take into account electrostatic interactions for calculating frustration, will be used. For more details, users can check our Frustratometer2 web server publication. Long range electrostatic interactions are modeled using a Debye-Hückel potential that includes both the solvent dielectric effect and the screening of charge-charge interactions by mobile ions in the solvent. By default a k = 4.15, corresponding to an aqueous solution, is assumed. The energy function used by frustratometeR considers Arg, Lys, Asp and Glu to be charged residues. Further information can be found in Tsai et al, Protein Science 2015.

SeqDist: Sequence separation used to calculate the local densities of the amino acids. This version implements by default a distance equal to 12 which is back compatible with the one used in the original frustratometeR algorithm (Jenik et. al, 2012). Additionally users can use a sequence distance equal to 3 which is in more agreement with results observed in molecular dynamics simulations using AWSEM-MD. Most published results in literature, using protein structures not in the context of molecular dynamics as produced by AWSEM-MD use *SeqDist=12*.

### Visualisations:

**The functions that generate graphs return ggplot2 objects modifiable to your liking!!**

##### Density of local frustration in a sphere of 5A for each residue of the structure.

The graph can be made in this way, taking the frustrated PDB object as a parameter. It can also be done for a specific chain.

```
plot_5Andens(Pdb_conf)
```

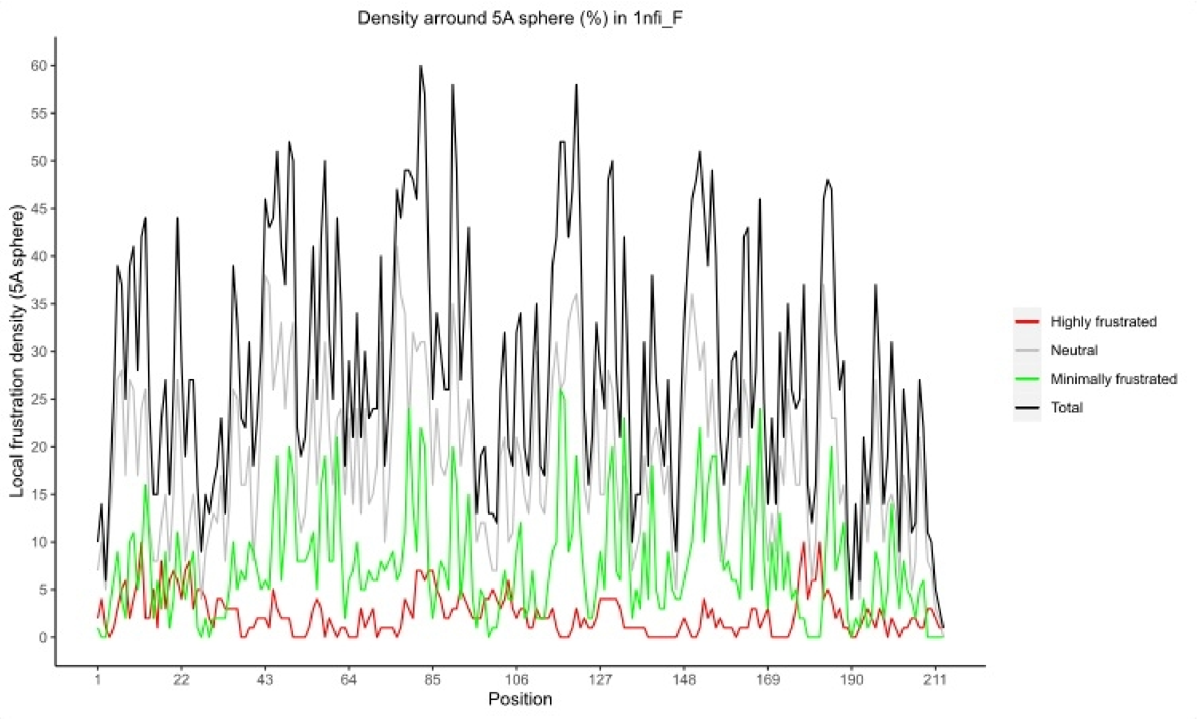


##### Proportion of density of local frustration in a sphere of 5A for each residue in the structure.

The graph can be made in this way, taking the frustrated PDB object as a parameter. It can also be done for a specific chain.

```
plot_5Adens_proportions(Pdb_conf)
```

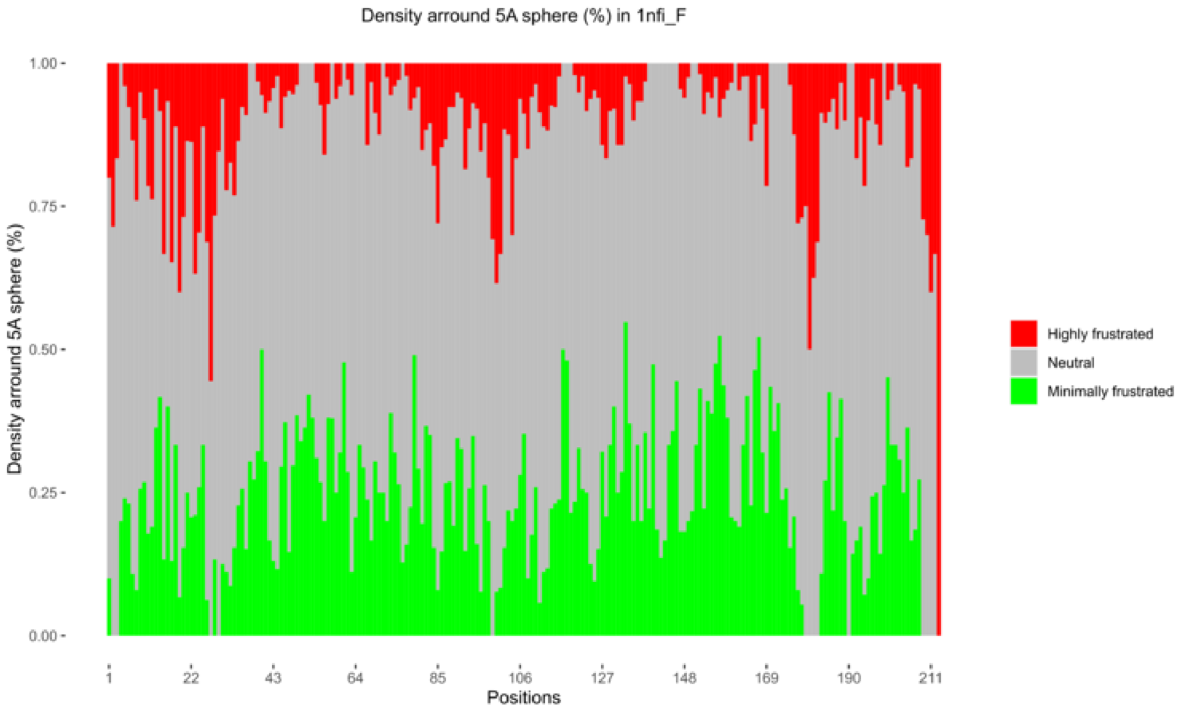

##### Contact map between structure residues with their respective frustration

The graph can be made in this way, taking the frustrated PDB object as a parameter. It can also be done for a specific chain.

```
plot_contact_map(Pdb_conf)
```

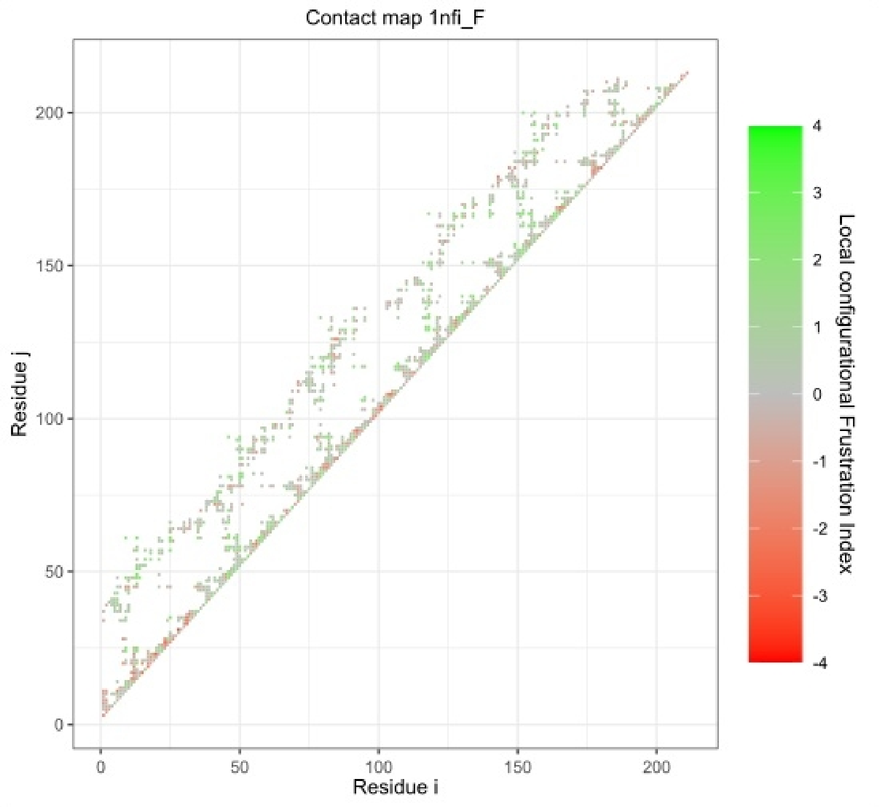


##### Pymol representation of the three-dimensional structure with its respective contacts.

The graph can be made in this way, taking the frustrated PDB object as a parameter. It can also be done for a specific chain. The contacts are represented with continuous lines (short or long) or dotted (water-mediated) and with different colours, in green (minimally frustrated) and red (highly frustrated). Neutral contacts are not shown (they are just too many!!).

```
view_frustration_pymol(Pdb_conf)
```

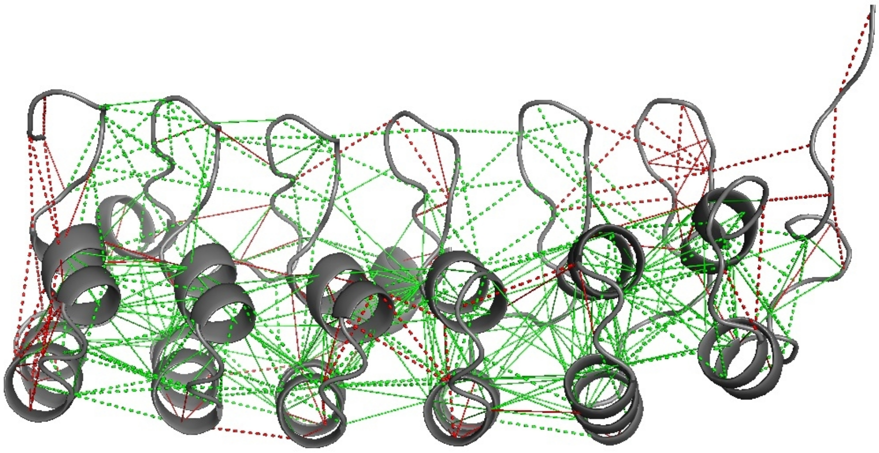

#### Summarising:

1. Have the structure in a file on your computer or know the *PdbId*.
2. Calculate the local energy frustration with the function `calculate_frustration()` obtaining a Pdb frustration object.
3. Get the results in table form for analysis with `get_frustration()`.
4. View the results:

> - 5Adens plot: `plot_5Adens()`
> - 5Adens proportion plot: `plot_5Adens_proportions()`
> - Contact map: `plot_contact_map()`
> - Pymol visualisation: `view_frustration_pymol()`

#### Full script:

#Load frustratometeR

```
library(frustratometeR)
```

#Calculate local energetic frustration

```
Pdb_conf <- calculate_frustration(PdbID = "1nfi", Chain = "F", Mode = "configurational")
```

#Obtain results for analysis

```
Frustration_data <- get_frustration(Pdb = Pdb_conf, Resno = 178, Chain = "F")
```

#Visualisations

```
plot_5Adens(Pdb_conf)
```

```
plot_5Adens_proportions(Pdb_conf)
```

```
plot_contact_map(Pdb_conf)
```

```
view_frustration_pymol(Pdb_conf)
```

### 4. Predicting frustration effect for mutations

#### Introduction the module to calculate changes in frustration because of mutations in the sequence/structure level.

The diagram shows that from the Pdb frustration object obtained by executing `calculate_frustration()`, explained in the previous entry, the 20 possible amino acid variants at a certain position can be analyzed. The `mutate_res()` function uses the frustration index previously specified in `calculate_frustration()`. This returns the modified Pdb frustration object, adding the data of the calculation performed. Thus, it can be used to obtain different visualisations.

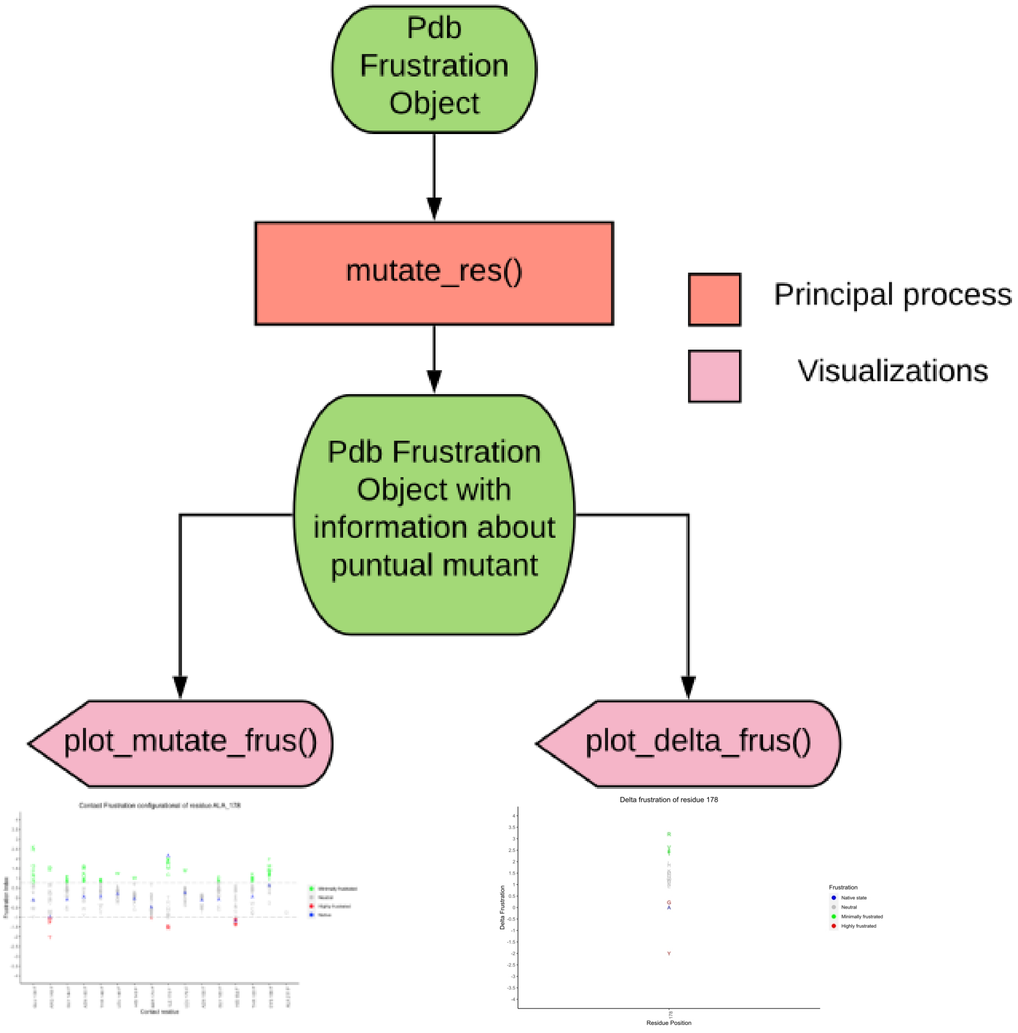


#### Example of analysis and explanation of use!

FrustratometeR provides us with a module that allows us to analyse the effect of point mutations on local frustration. It can be applied to single chains or protein complexes (quaternary structures) with multiple chains. The mutation effect can be evaluated for any of the 3 frustration indexes, i.e, "configurational", "mutational" or "singleresidue".

After using the `calculate_frustration()` function with one of the 3 frustration indexes, the `mutate_res()` function works in two steps as follows: 1) The input protein is mutated at a given position from the native amino acid to all the other 19 amino acids. This is made by using one of two modes:

- **Threading**: It will change the amino acid identity (it will introduce a different side chain to the structure) but it will not modify the rest of the protein backbone.
- **Modeller**: It will change the amino acid identity and on top of it, it will perform an homology model with a small energetic optimisation of the structure. This may be reflected in structural variations in the coordinates from other regions of the protein, beyond the mutated amino acid. This mode is slower than the threading one.

2) Frustration values, using the same mode that was used with the `calculate_frustration()` function, are calculated for the new "mutated" structures and stored in the R object(Pdb frustration object) to be visualised or retrieved by other functions.

The `mutate_res()` function can receive a parameter called *Split*. In case of a multimeric protein structures users can decide if after the mutation is made frustration should be calculated to a specific chain or the entire complex by specifying using the *Split*, parameter. *Split* = TRUE specific chain, *Split* = FALSE full complex. This is important because frustration values will vary depending on using a quaternary structure or isolated chains. This is because the interacting partner offers extra contacts that change the local energetics.

Minum steps:

**1st step:** calculate frustration for the native structure.

```
Pdb_conf <- calculate_frustration(PdbID = "1nfi", Chain = "F", Mode = "configurational")
```

**2nd step:** mutate a given residue.

This will first mutate the residue using the threading mode by default (*Method*="Threading") or alternatively using the Modeller mode (*Method*="Modeller").

```
Pdb_conf <- mutate_res(Pdb = Pdb_conf, Resno = 178, Chain = "F")
```

### Visualisations:

**The functions that generates graphs return ggplot2 objects modifiable to your liking!!**

The configurational and mutational modes are contact specific. Every time we mutate a residue, we can change the frustration values for the native contacts, some contacts may disappear and new ones can be established as well. To summarise what happens when we mutate a certain residue by all alternative aminoacids we generate specific visualisations. For this, the function receives the object of frustration Pdb, the residue number, chain structure is located and the mutation method.

```
plot_mutate_res(Pdb = Pdb_conf, Resno = 178, Chain = "F")
```

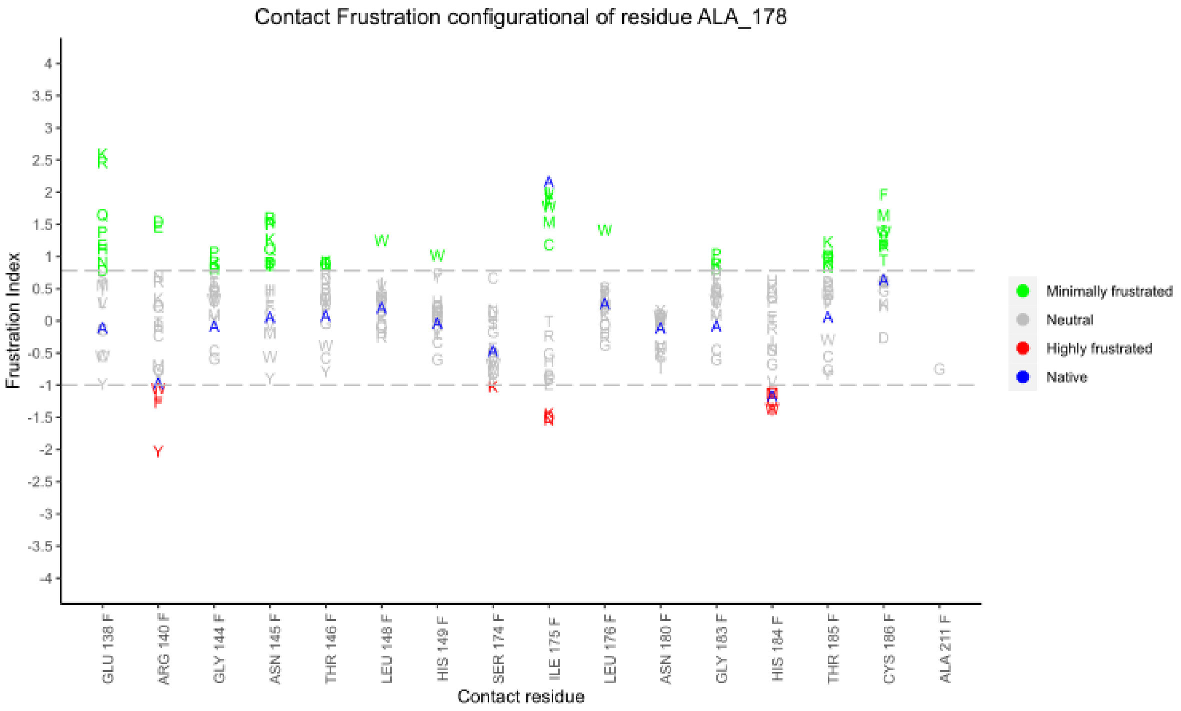


For example, in the graph it can be seen that Ala 178 allows the most favorable interaction with residue Ile 175 of the F chain when comparing with each of the 19 amino acid variants. Meanwhile, the interaction with the Ala 211 residue of the F chain is only formed in the case that there is a Gly at position 178.

For the single residue mode, since it is not by contact, we can make a graph that shows the difference in frustration with respect to the native (in structure). This requires indicate Pdb frustration object obtained from the execution of `calculate_frustration()` with *Mode = "singleresidue"* and `mutate_res()` for the specific residue. Also, the residue number, Chain where it is found and the mutation method.

```
plot_delta_frus(Pdb = Pdb_sing, Resno = 178, Chain = "F")
```

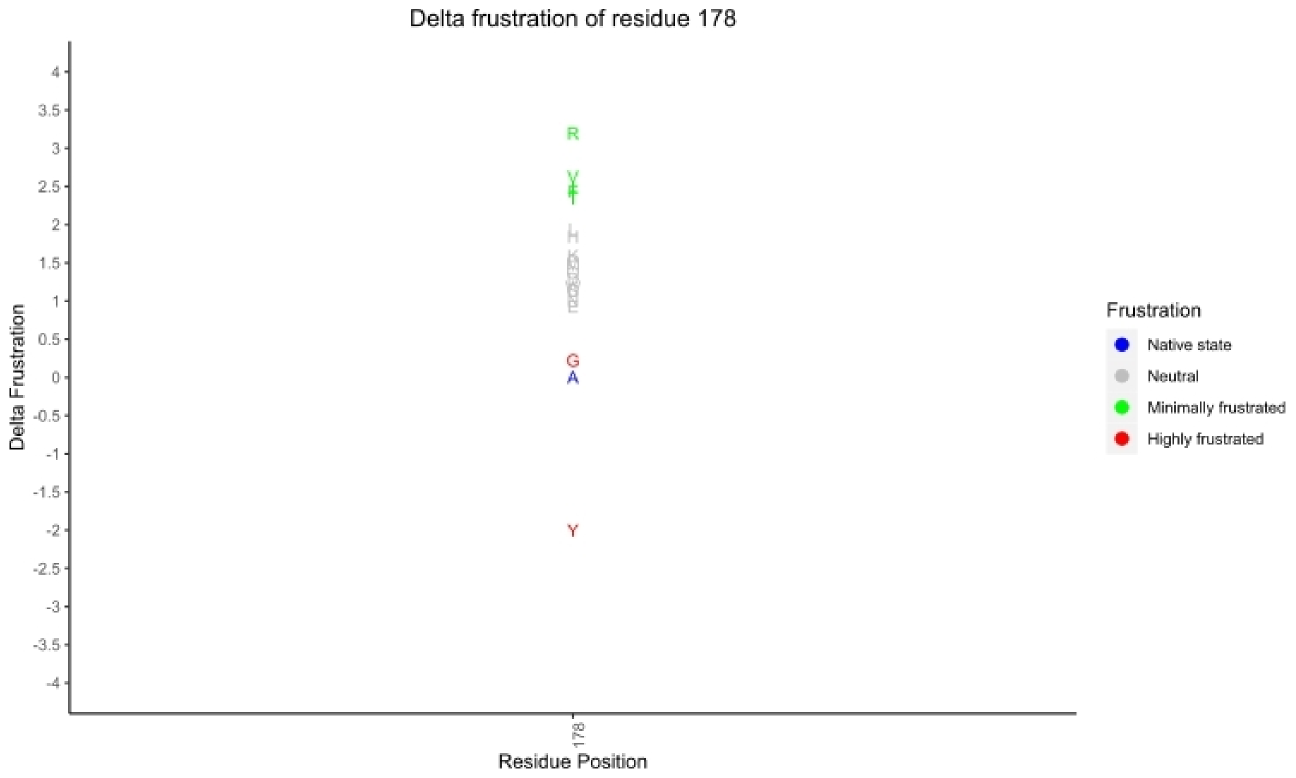


#### Summarising

The steps to predict the effect of point mutations on local frustration are:

1. Get Pdb frustration object with `calculate_frustration()`. It should be noted that the desired frustration index must be selected here.
2. Mutate a specific residue of the structure with `mutate_res()`, obtaining the modified Pdb frustration object.
3. Use the Pdb frustration object to obtain graphics with `plot_mutate_res()` and `plot_delta_frus()`, depending on the frustration index used.

#### Full script

#Load frustratometeR

```
library(frustratometeR)
```

##### With frustration index at the contact level (configurational or mutational)

#Obtain Pdb frustration object

```
Pdb_conf <- calculate_frustration(PdbID = "1nfi", Chain = "F", Mode = "configurational")
```

#Calculate local frustration for 20 amino acid variants for residue 178 in chain F

```
Pdb_conf <- mutate_res(Pdb = Pdb_conf, Resno = 178, Chain = "F")
```

#View results

```
plot_mutate_res(Pdb = Pdb_conf, Resno = 178, Chain = "F")
```

##### With frustration index at the single-residue level

#Obtain Pdb frustration object

```
Pdb_sing <- calculate_frustration(PdbID = "1nfi", Chain = "F", Mode = "singleresidue")
```

#Calculate local frustration for 20 amino acid variants for residue 178 in chain F

```
Pdb_sing <- mutate_res(Pdb = Pdb_sing, Resno = 178, Chain = "F")
```

#View results

```
plot_delta_frus(Pdb = Pdb_sing, Resno = 178, Chain = "F")
```

### 5. Molecular dynamics analysis

#### Introduction to the local energy frustration analysis module in molecular dynamics simulations

The diagram below shows how, from a directory containing the structures of each frame of a molecular dynamics, the local energy frustration can be analyzed with `dynamic_frustration()`. Generates a dynamic frustration object that can be used to get visualisations and other processing.

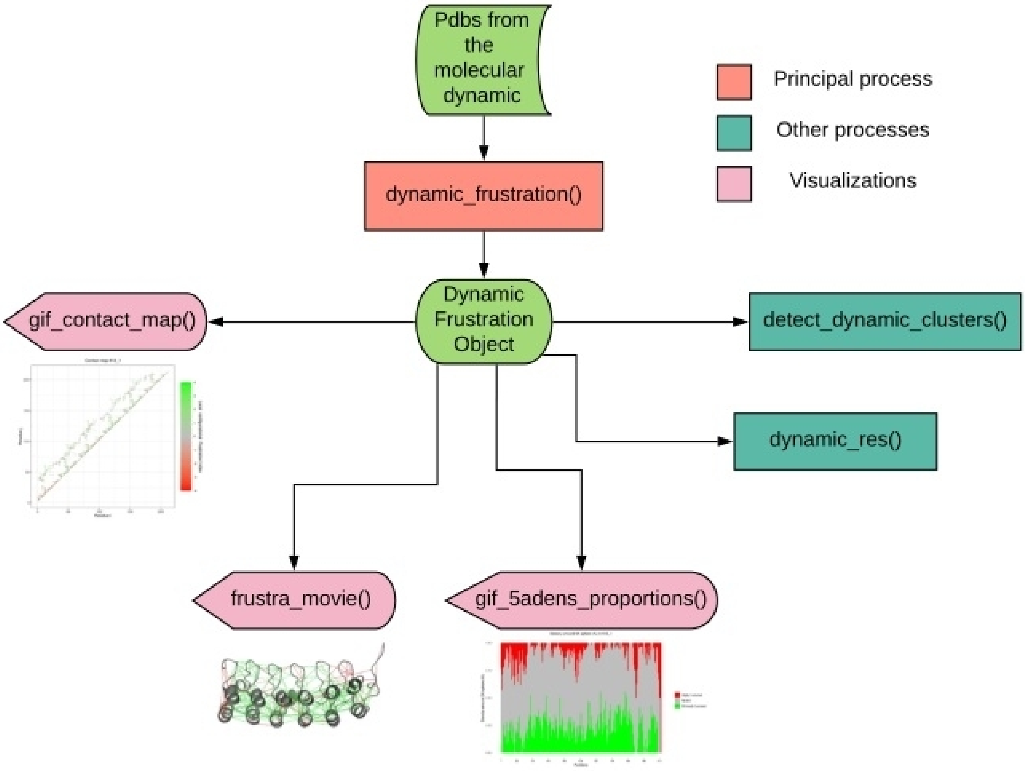


The diagram below shows one of the possible processes of a dynamic frustration object obtained with `dynamic_frustration()`. With the `dynamic_res()` function you can analyze the frustration dynamics of a certain residue. This returns a modified dynamic frustration object, adding the resulting data from the process. The dynamic frustration object can be used to get the frustration data or make visualisations.

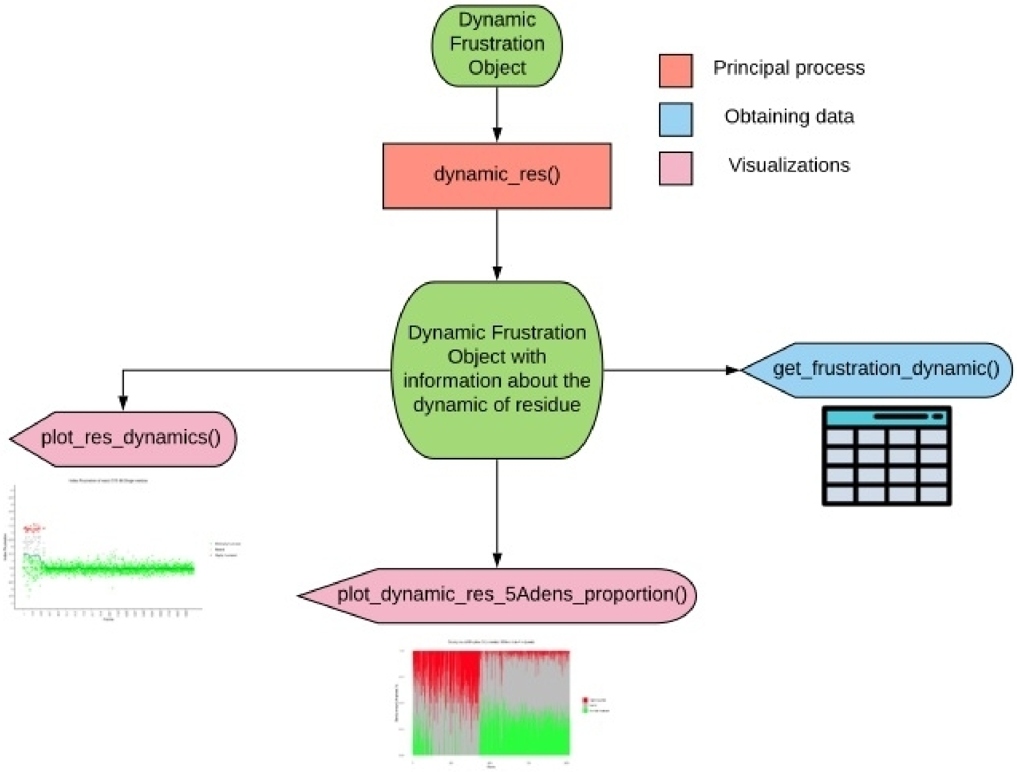

### Example of analysis and explanation of use!

FrustratometeR allows us to analyse frustration along a molecular dynamics trajectory. In this case we are going to explain these new functionalities by analysing a molecular dynamics trajectory obtained for the IkappaB-alpha protein (PDBID: 1nfi,F). The trajectory was generated by using an coarsed grained AMH-GO folding model as implemented in the AWSEM-MD suit (Davtyan et al, 2012). After estimating the folding temperature (Tf), we applied an umbrella sampling method to simulate the dynamics of the protein along the Qw coordinate (Qw corresponds to the fraction of native contacts in a given structure compared to the initial structure being used). The Qw coordinate, that can take values between 0 (completely unfolded) and 1 (completely folded) was sampled by dividing the interval in 40 bins, simulating 10 millions steps every 3 femtoseconds. All 40 simulations are finally integrating using WHAM (Weighted Histogram Analysis Method) obtaining Free Energy values and projections of those into different coordinates like Qw or the Radius of Gyration (Rg). More information on how to perform this type of simulations can be found in Schaefer et al, 2014.

Below you can find a Free energy diagram projected into the Qw and Rg coordinates. Different sampled conformations are shown for different characteristic regions in the landscape. U: Unfolded state, F: Folded State, I: Folding intermediary, E1: Expanded state 1, E2: Expanded state 2.

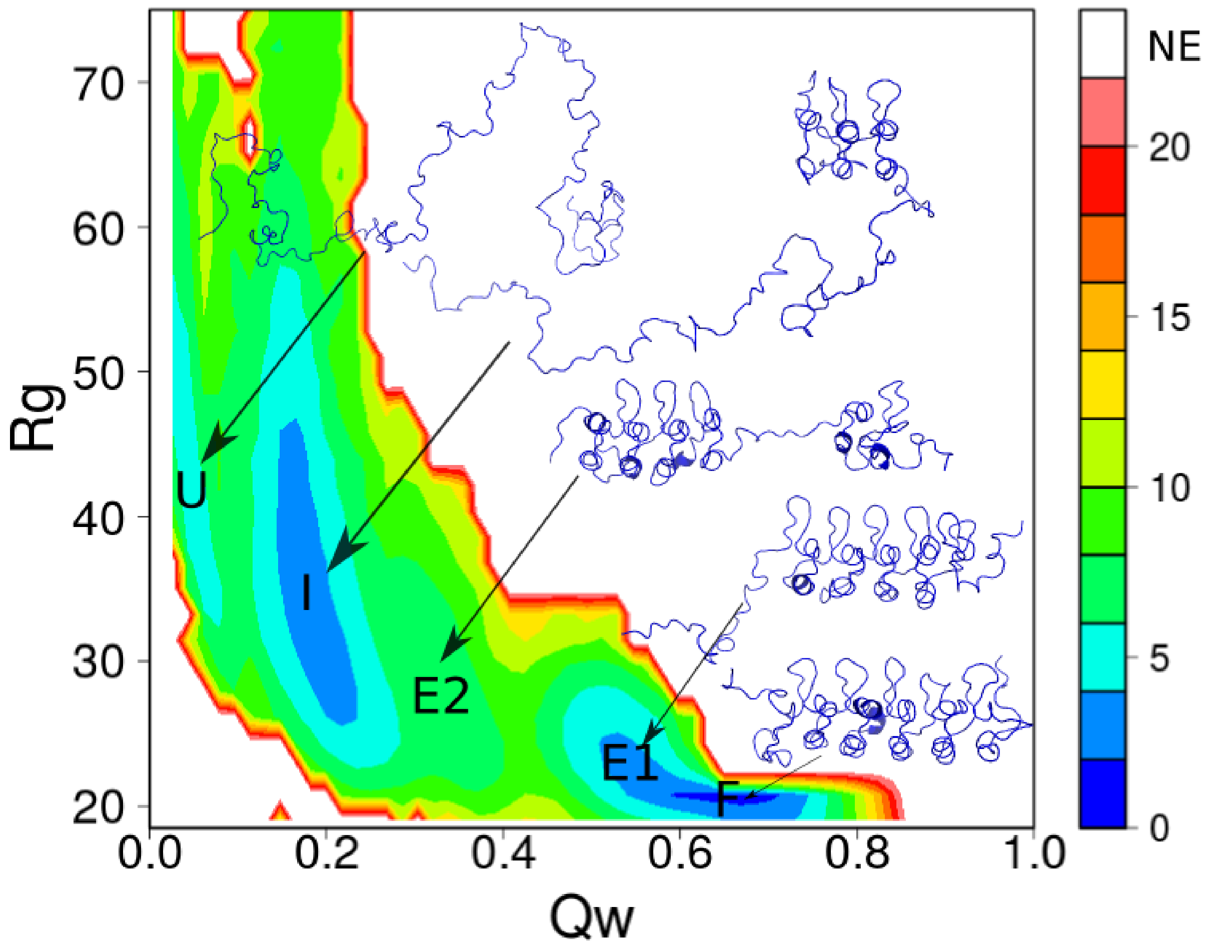


From this full folding trajectory, we subsampled 2040 equidistant frames. These frames can be downloaded from https://github.com/proteinphysiologylab/frustratometer-data.git and placed into a specific folder in your computer.

A vector must be created that contains the names of the structures in the desired temporal order:

```
OrderList <- paste0("612_", 1:2040, ".pdb")
```

In this example we have 2040 structures called pdb [i] .pdb, where i goes from 1 to 2040.

We use the `dynamic_frustration()` function to analyse the trajectory. The full path of the directory where the structures are located (*PdbsDir*), the order in which they will be analysed (*OrderList*, optional) and the desired frustration index (*Mode*) have to be passed as parameters:

```
Dynamic_conf <- dynamic_frustration(PdbsDir = "/home/Dynamic/pdbs/", OrderList = OrderList, ResultsDir = "/home/Dynamic/results/", Mode = "configurational")
```

The previous step can take a long time to run, depending on the size of the protein and the number of provided frames. Users can save the object in case they want to analyse it at later times:

```
saveRDS(Dynamic_conf, "personalfolder/dynamic_object.RDS")
```

Once finished the dynamic frustration object can be parsed to extract the local frustration profile for a specific residue by using the `dynamic_res()` function. We must indicate the Dynamic frustration object (*Dynamic*), residue number (*Resno*) and chain (*Chain*).

```
# The chain parameter needs to be specified since the provided pdb structure could be a multimeric complex.
```

```
Dynamic_conf <- dynamic_res(Dynamic = Dynamic_conf, Resno = 185, Chain = "A")
```

The dynamic information for the specified residue will be stored in the Dynamic frustration object (Dynamic_conf).

The energetic local frustration results can be obtained in the form of a Data Frame with the `get_frustration_dynamic()` function. You can specify a specific residue (*Resno*) or chain (*Chain*), as well as certain frames listed in vector form ( *Frames*):

```
DynamicFrusData <- get_frustration_dynamic(Dynamic = Dynamic_conf, Resno = 185, Chain = "A", Frames = c(1, 5, 10))
```

In this case, a table is obtained containing the local energetic frustration of residue 185 of chain A in frames 1, 5 and 10 of the *Dynamic_conf* dynamic.

#### Visualisations:

Results can be visualised with different functions in the same way as for individual structures.

Local frustration proportions (sphere 5A) for each residue:

```
gif_5adens_proportions(Dynamic = Dynamic_conf)
```

Contact map:

```
gif_contact_map(Dynamic = Dynamic_conf)
```

Also, a Frustration Movie showing the frustration patterns projected on top of the folding three-dimensional structure can be generated. To do this initially, the script [*ResultsDir*] /FrustraMovie/representations_configurational.pml must be opened with Pymol, which will load each frame to adjust the viewing angle. When you have chosen it, run the script [*ResultsDir*] /FrustraMovie/GenerateMovie_configurational.pml which will create the movie that you can store as a gif.

```
frustra_movie(Dynamic = Dynamic_conf)
```

Static plots will produce different results depending on which frustration index was selected when the object was created.

**The gifs can be viewed at this link!**

For the configurational or mutational indexes:

```
plot_res_dynamics(Dynamic = Dynamic_conf, Resno = 185, Chain = "A")
```

```
plot_dynamic_res_5Adens_proportion(Dynamic = Dynamic_conf, Resno = 185, Chain = "A")
```

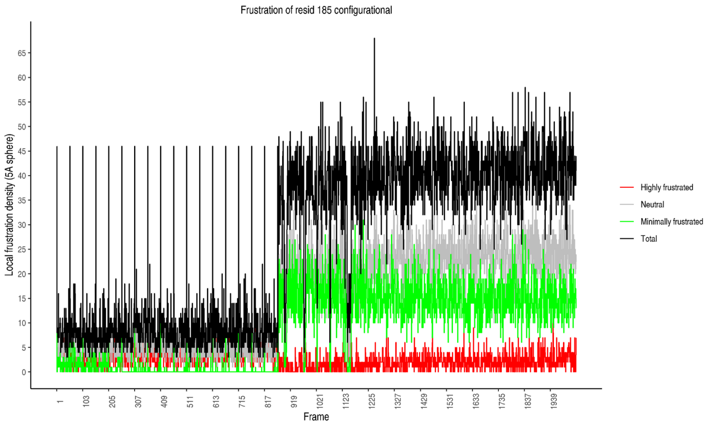


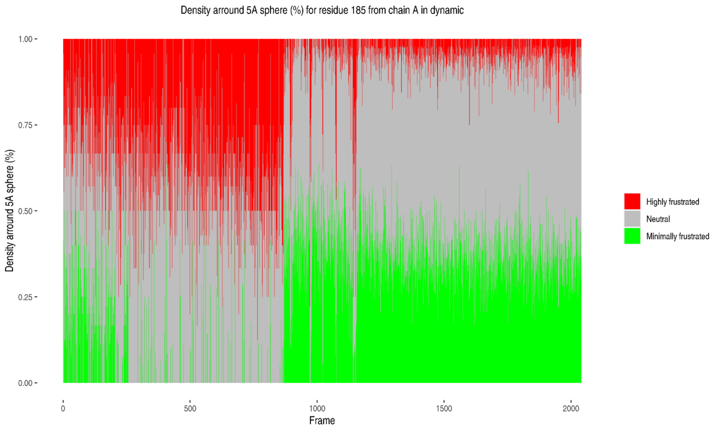

For the singleresidue index:

```
plot_res_dynamics(Dynamic = Dynamic_sing, Resno = 185, Chain = "A")
```

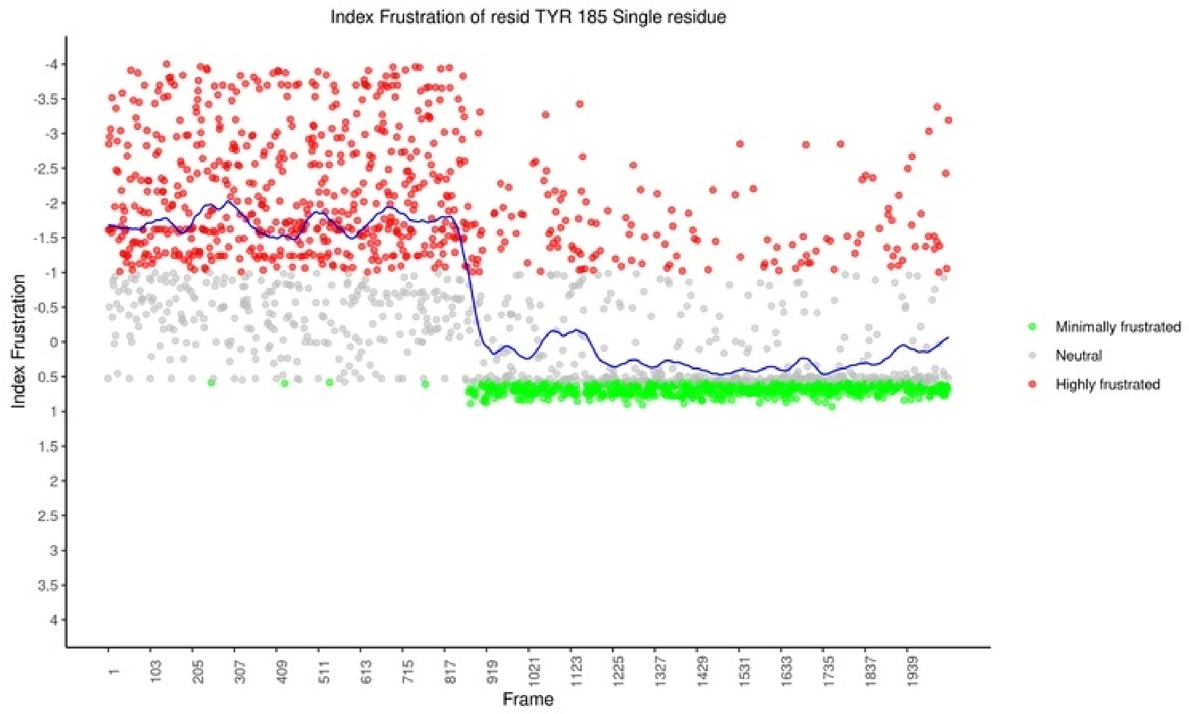


#### Summarising

The steps to analyse local frustration throughout a dynamic are:

1. Contain in a directory the structures that represent each frame of the dynamics.
2. Calculate the local frustration of all or part of the dynamics with `dynamic_frustration()`.
3. Analyse local frustration in the dynamics of a specific residue with `dynamic_res()`.
4. Get the results in table form for analysis with `get_frustration_dynamic()`.
5. Obtain the necessary graphics:
    - Gifs: `gif_5adens_proportions()` and `gif_contact_map()`.
    - Graphics: `plot_res_dinamics()` and `plot_dynamic_res_5Adens_proportion()`.
    - FrustraMovie: `frustra_movie()`

#### Full script

#Load frustratometeR

```
library(frustratometeR)
```

### Defining the order of the structures. We create a vector with the names of the pdb file in the order that we want FrustratometeR to process them

```
OrderList <- paste0("612_", 1:2040, ".pdb")
```

##### With frustration index at the contact level (configurational or mutational)

#Calculation of local energy frustration for all frames of dynamics

```
Dynamic_conf <- dynamic_frustration(PdbsDir = "/home/Dynamic/pdbs/", OrderList = OrderList, ResultsDir = "/home/Dynamic/results/", Mode = "configurational")
```

#Local frustration along the dynamics for a specific residue

```
Dynamic_conf <- dynamic_res(Dynamic = Dynamic_conf, Resno = 185, Chain = "A")
```

#Obtain the results for residue 185 of chain A for frames 1, 5 and 10

```
DynamicFrusData <- get_frustration_dynamic(Dynamic = Dynamic_conf, Resno = 185, Chain = "A", Frames = c(1, 5, 10))
```

#Visualisations

```
plot_res_dynamics(Dynamic = Dynamic_conf, Resno = 185, Chain = "A")
```

```
plot_dynamic_res_5Adens_proportion(Dynamic = Dynamic_conf, Resno = 185, Chain = "A")
```

#### With frustration index at the single-residue level

#Calculation of local energy frustration for all frames of dynamics

```
Dynamic_sing <- dynamic_frustration(PdbsDir = "/home/Dynamic/pdbs/", OrderList = OrderList, ResultsDir = "/home/Dynamic/results/", Mode = "singleresidue")
```

#Local frustration along the dynamics for a specific residue

```
Dynamic_sing <- dynamic_res(Dynamic = Dynamic_sing, Resno = 185, Chain = "A")
```

#Visualisation

```
plot_res_dynamics(Dynamic = Dynamic_sing, Resno = 185, Chain = "A")
```

#### Obtaining gifs and FrustraMovie

The same dynamics were used but only 50 frames were taken from 1020 to 1070 due to computational cost. Thus:

```
OrderList <- paste0("612_", 1020:2070, ".pdb")
```

#Calculation of local energy frustration for all frames of dynamics

```
Dynamic_conf <- dynamic_frustration(PdbsDir = "/home/Dynamic/pdbs/", OrderList = OrderList, ResultsDir = "/home/Dynamic/results/", Mode = "configurational")
```

#Gifs and FrustraMovie

```
gif_5adens_proportions(Dynamic = Dynamic_conf)
```

```
gif_contact_map(Dynamic = Dynamic_conf)
```

```
frustra_movie(Dynamic = Dynamic_conf)
```

### 6. Detect dynamic clusters

#### Introduction to detection of dynamic clusters

The following diagram shows one of the processes that can be applied to the Pdb dynamic object obtained with `dynamic_frustration()`. This process is `detect_dynamic_clusters()` which finds clusters of residues with similar frustration dynamics. It modifies the Pdb dynamic object that can be used to obtain the information of the clusters and multiple visualisations.

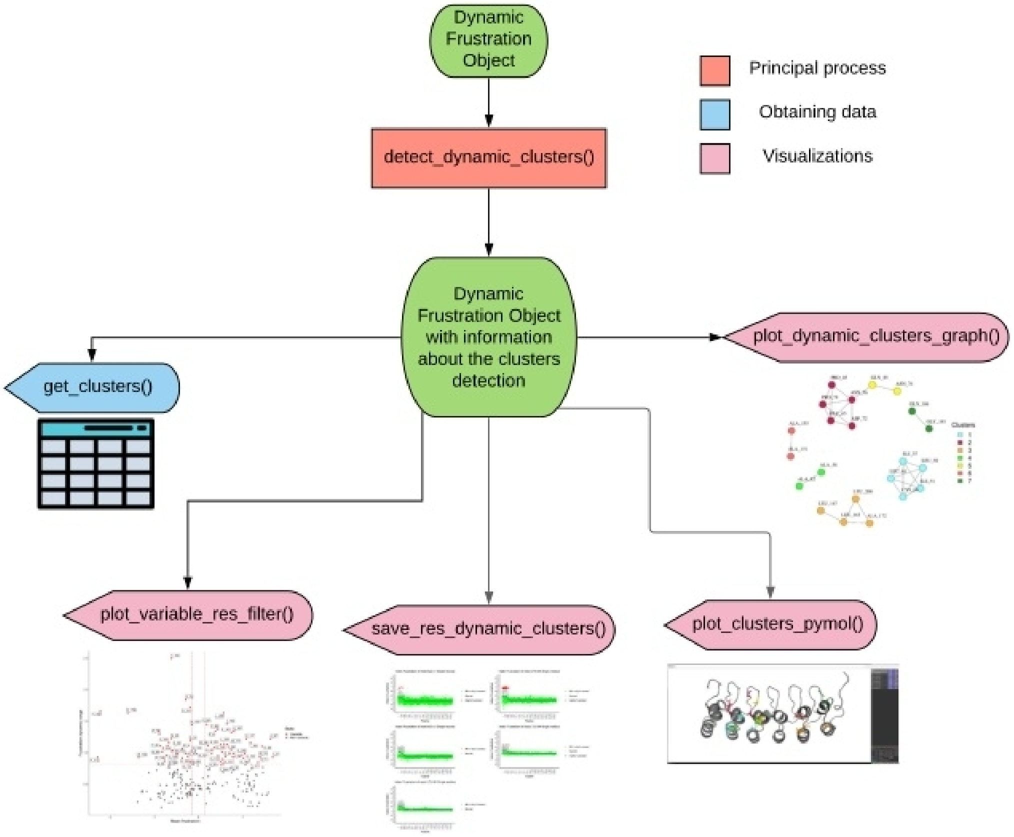


##### This example shows the use of dynamic clusters discovery and interpretation

FrustratometeR has the additional function of detecting residues that have correlated changes in their local energetic frustration and variants throughout a molecular dynamics trajectory.

In this case we are going to explain these new functionalities by analysing a molecular dynamics trajectory obtained for the IkappaB-alpha protein (PDBID: 1nfi,F) that we started analysing in the previous entry.

From this full folding trajectory, we subsampled 2040 equidistant frames. These frames can be downloaded from https://github.com/proteinphysiologylab/frustratometer-data.git and placed into a specific folder in your computer.

In order to detect dynamic clusters, the singleresidue frustration index is used (*Mode = "singleresidue"*).

**1st step:** Calculate frustration values for all the trajectory frames using the `dynamic_frustration()` function:

```
OrderList <- paste0("612_", 1:2040, ".pdb")
```

```
Dynamic_sing <- dynamic_frustration(PdbsDir = "/Folder/IkappaBalfa_dynamic_2040Frames/", OrderList = OrderList, ResultsDir = "/Folder/IkappaBalfa_dynamic_2040Frames/results/", Mode = "singleresidue")
```

**2nd step:** Detect dynamic clusters using the `detect_dynamic_clusters()` function. Be aware that in what follows we have tuned the default parameters so the function will produce meaningful results in the context of our system. In general terms you can always start with the default parameters and further modify them to assess how much sense they make.

```
Dynamic_sing <- detect_dynamic_clusters(Dynamic = Dynamic_sing, CorrType = "spearman", FiltMean = 0.15, MinFrstRange = 0.7, MinCorr = 0.95)
```

`detect_dynamic_clusters()` filters the protein residues based on two parameters:

1) **"Frustration mean"** across the trajectory: The *FiltMean* (Mean filter threshold) is used to filter out those residues with a mean absolute value across the dynamics lower than *FiltMean*.

2) **"Frustration dynamic range"**: Only those residues with a Frustration dynamic range higher than *MinFrstRange* will be kept. A piecewise linear model is fitted to the frustration dynamic profiles (*LoessSpan*: controls the degree of smoothing) for all residues. The linear model is used to calculate the frustration range of variation across the trajectory (the difference between the maximum and minimum point in the adjusted model).

Residues that pass these two filters will be kept to build the clusters. A Principal Component Analysis (PCA) is calculated and the *Ncp* parameter is used to define how many components are kept (default value *Ncp*=10). Correlation values are calculated (either using "spearman" by default or "pearson") in the PCA space between all pairs of residues. The *MinCorr* parameter defines which correlation coefficients will be kept to define that two residues are connected in the network. The resulting adjacency matrix is used to generate an undirected graph and the Leiden clustering method (Traag et al, 2019) is used to detect the clusters. The *LeidenResol* parameter defines the resolution at which clusters will be detected (default value 1.00).

**Note: All these parameters have to be explored and adjusted for each specific system in order to produce meaningful results.**

Clusters are stored in the Dynamic frustration object. Cluster assignments can be retrieved by:

`get_clusters(Dynamic_sing, Clusters = "all")`.

Specific clusters can be retrieved by changing the *Clusters* parameter as:

`get_clusters(Dynamic_sing, Clusters = c(1,2))`

#### Visualisations:

A plot showing the spread of residues in terms of their average frustration values and frustration dynamic range can be visualised with:

`plot_variable_res_filter(Dynamic = Dynamic_sing)`

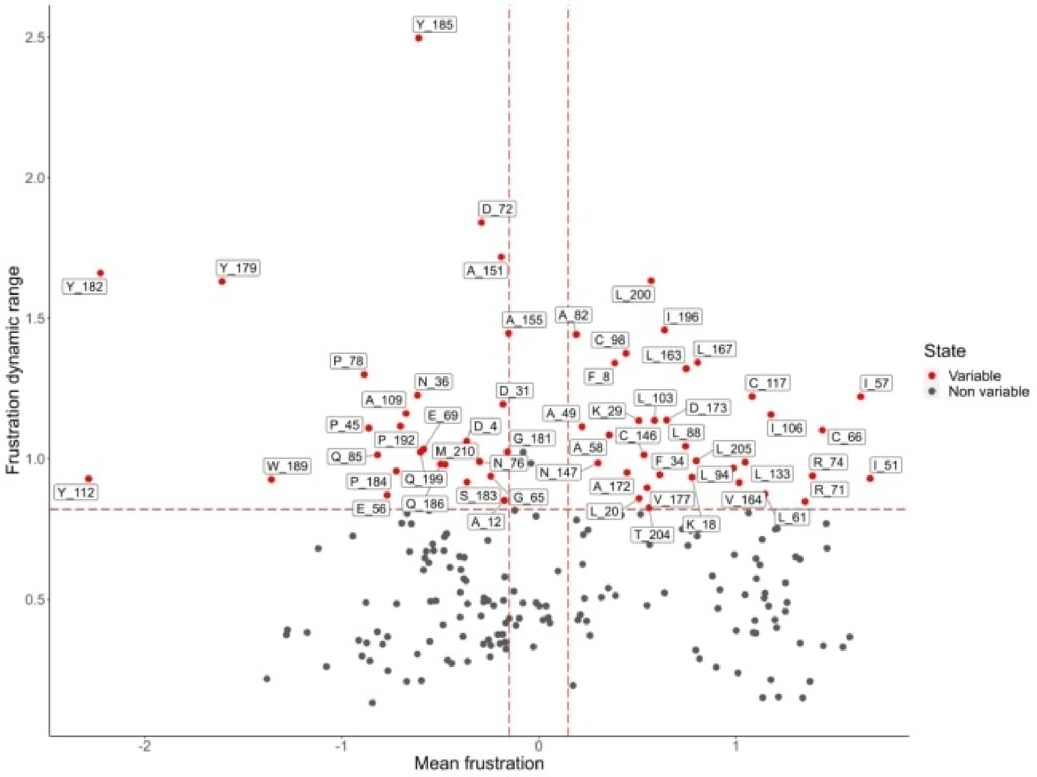

Residues that passed the filters in the previous plot and that are clustered using the Leiden algorithm can be visualised with:

```
plot_dynamic_clusters_graph(Dynamic = Dynamic_sing)
```

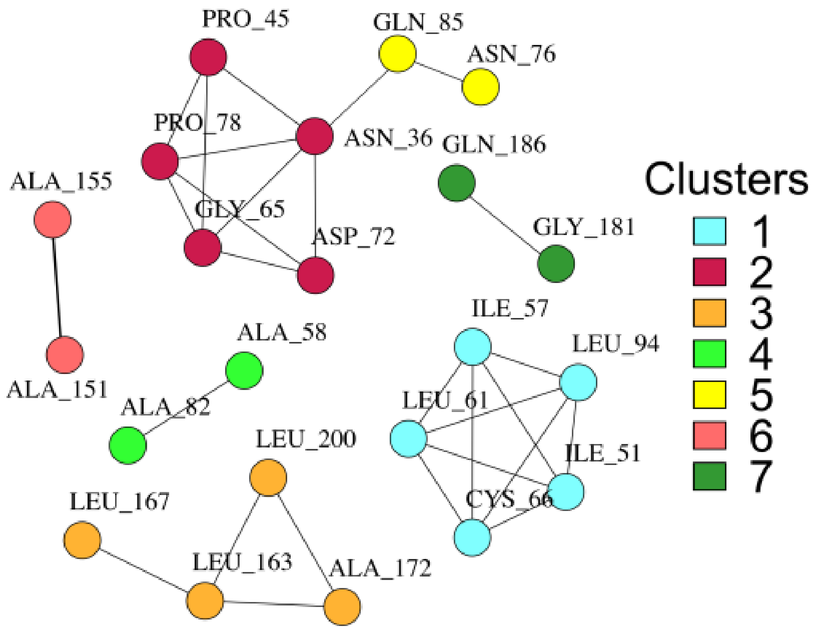


Note that in the undirected graph there are fewer residuals than passed the filter in frustration mean and frustration dynamic range. This is due to the fact that the adjacency matrix was formed with the residuals that have a correlation greater than *MinCorr=0.95*.

These residues can also be interactively visualised on top of the protein structure using Pymol with (the same colour scheme as in the previous plot is used to identify clusters):

```
plot_clusters_pymol(Dynamic_sing, Clusters = "all")
```

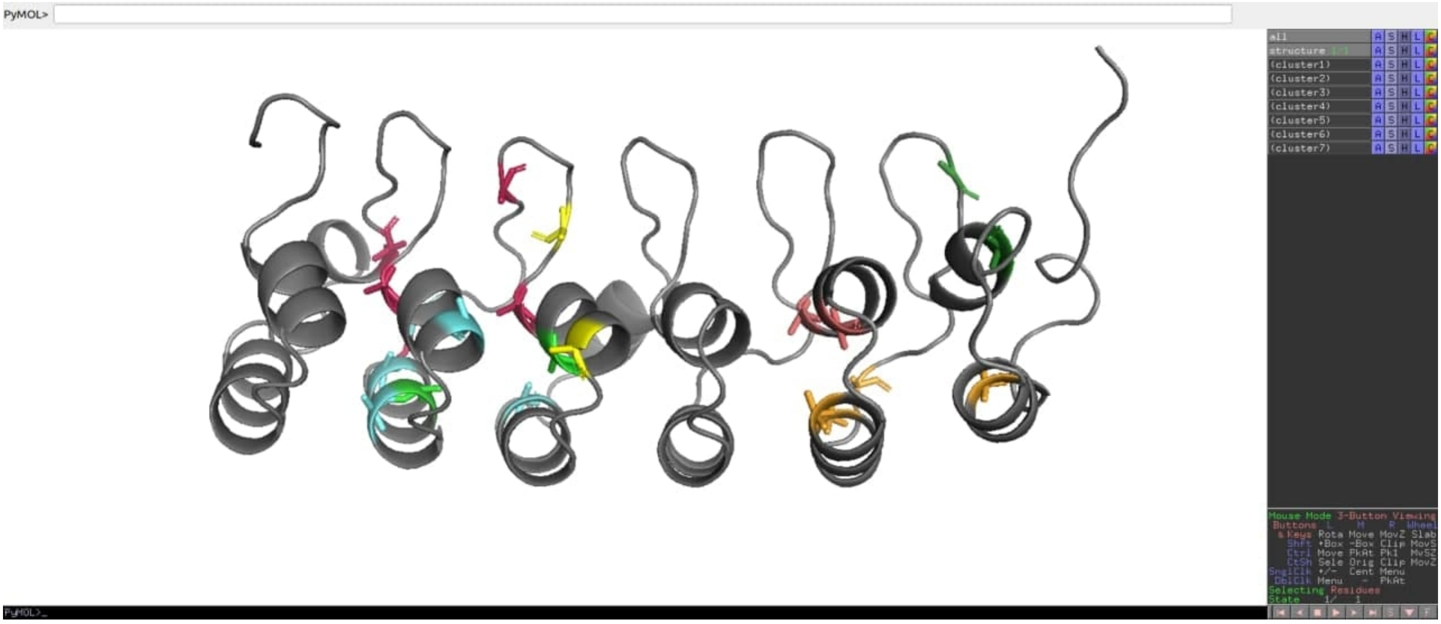


Frustration profiles for specific residues across the entire trajectory can be visualised with:

```
plot_res_dynamics(Dynamic = Dynamic_sing, Resno = 85, Chain = "A")
```

```
plot_res_dynamics(Dynamic = Dynamic_sing, Resno = 167, Chain = "A")
```

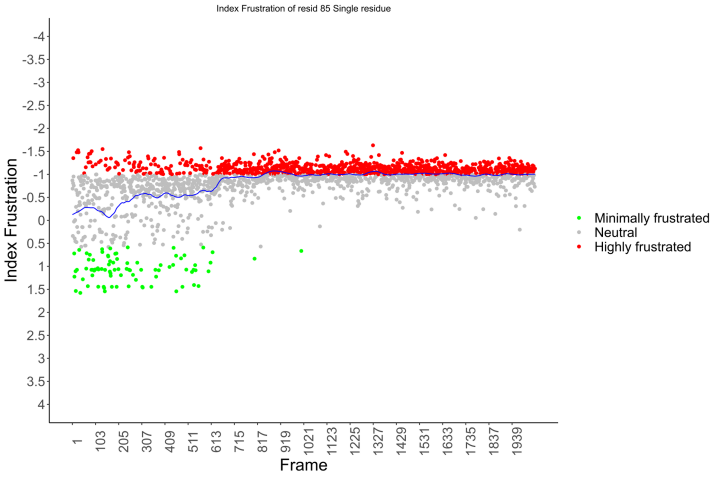


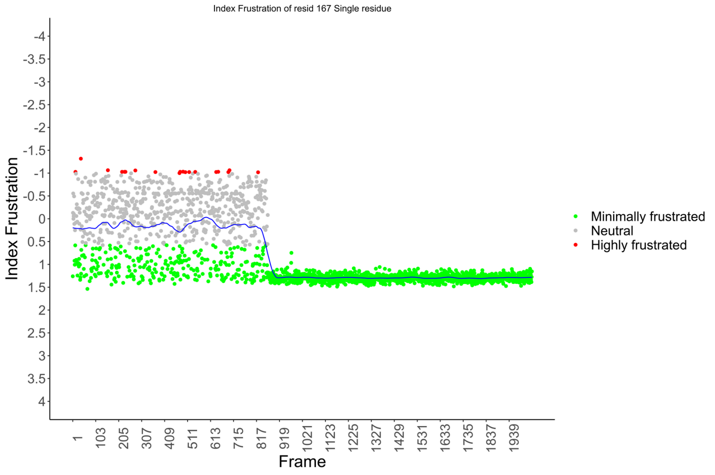


##### Interpreting the biological meaning of the clusters for IkBa (IkappaBAlpha) in the context of recognising NFkB:

We provide here more information about how the simulation was performed and how to interpret the results in order to assess the meaning for the clusters that were found. The trajectory was generated by using an coarsed grained AMH-GO folding model as implemented in the AWSEM-MD suite (Davtyan et al, 2012). After estimating the folding temperature (Tf), we applied an umbrella sampling method to simulate the dynamics of the protein along the Qw coordinate (Qw corresponds to the fraction of native contacts in a given structure compared to the initial structure being used). The Qw coordinate, that can take values between 0 (completely unfolded) and 1 (completely folded) was sampled by dividing the interval in 40 bins, simulating 10 millions steps every 3 femtoseconds. All 40 simulations are finally integrated using WHAM (Weighted Histogram Analysis Method) obtaining Free Energy values and projections of those into different coordinates like Qw or the Radius of Gyration (Rg). More information on how to perform this type of simulations can be found in Schaefer et al, 2014. Below you can find a Free energy diagram projected into the Qw and Rg coordinates. Different sampled conformations are shown for different characteristic regions in the landscape. **U:** Unfolded state, **F:** Folded State, **I:** Folding intermediary, **E1:** Expanded state 1, **E2:** Expanded state 2.

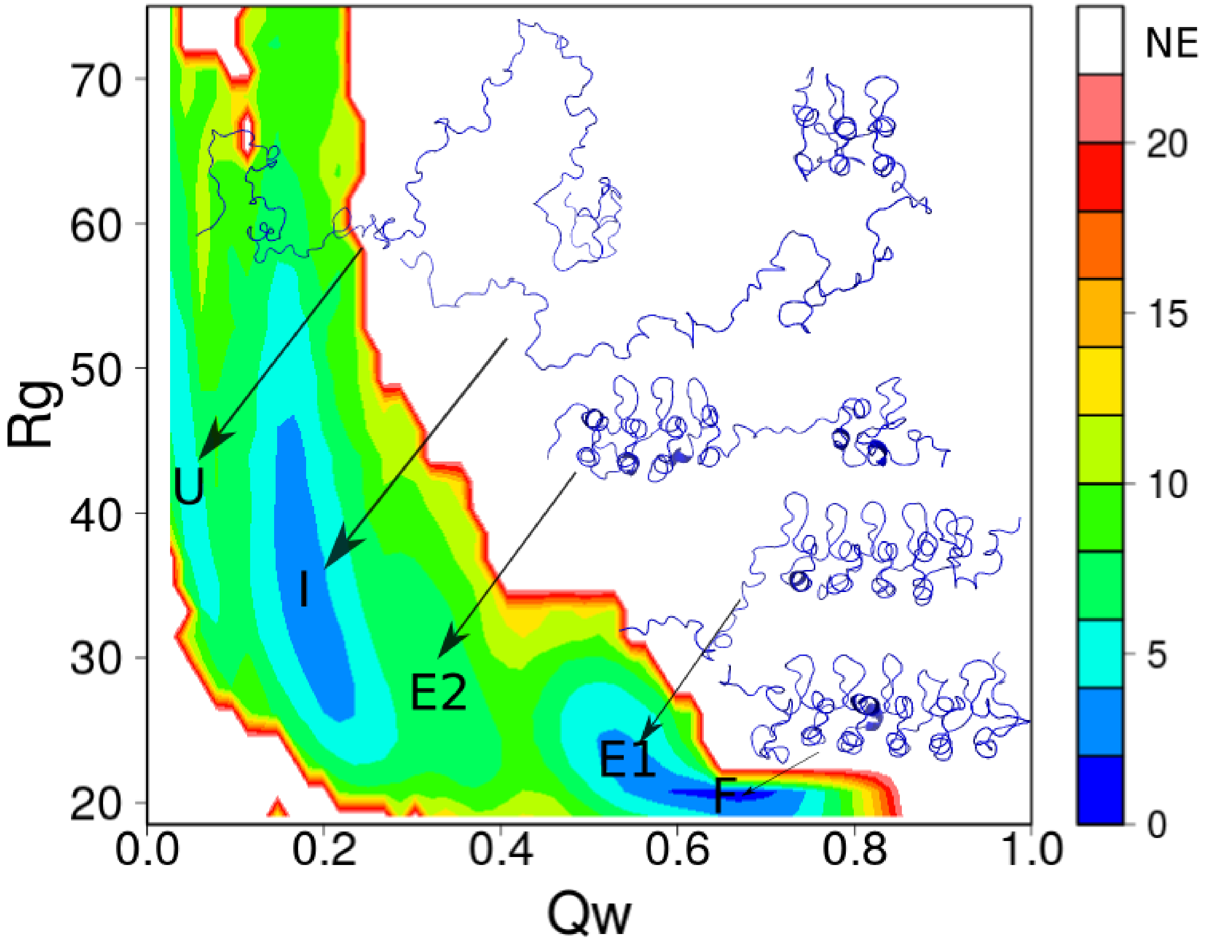


To understand the meaning of the clusters that we've found, we have to analyse the residues frustration profiles in the context of IkBa energy landscape. Remember also that IkBa folding is coupled to NFkB recognition (check Ferreiro & Komives, Biochemistry 2010). **The IkappaB/NfKB complex exhibits a very complex dynamic and it is not the aim of this tutorial to fully explain it but to offer some qualitative insights about how frustratometeR can be of help to analyse and further understand folding dynamics.**

**1st Folding Transition:**

IkBa has a 1st folding transition when the 1st two repeats start to fold, coupled to a first interaction with NfKB. The transition corresponds to the barrier that exists between the Unfolded state (U) and the Folding Intermediate state (I). Clusters 1, 2 and 4 correspond to residues that change their frustration state at around that barrier. Cluster 1 and Cluster 4 correspond to residues that become minimally frustrated and are located at Repeats1 and 2, mainly at the helices and are important for their stability. Cluster 2 corresponds to residues that become highly frustrated during this folding transition because they are important for the interaction with NFkB. The reason why they become highly frustrated is because NFKappaB is not present in the simulation (remember that is a AMH GO simulation where IkBa is artificially separated from NFkB but the structure is still forced to adopt the conformation that corresponds to the complex with NfKB) to energetically compensate their interactions. In fact not having NFkB is what allows us to further understand IkBa folding mechanism with an AMH GO model.

#### Cluster 1

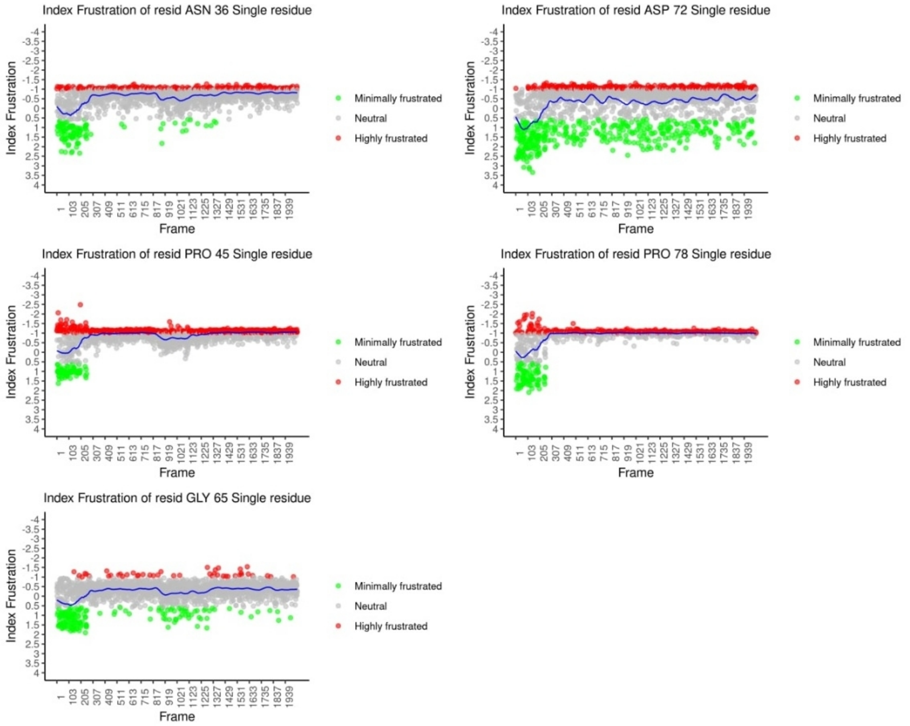


#### Cluster 2

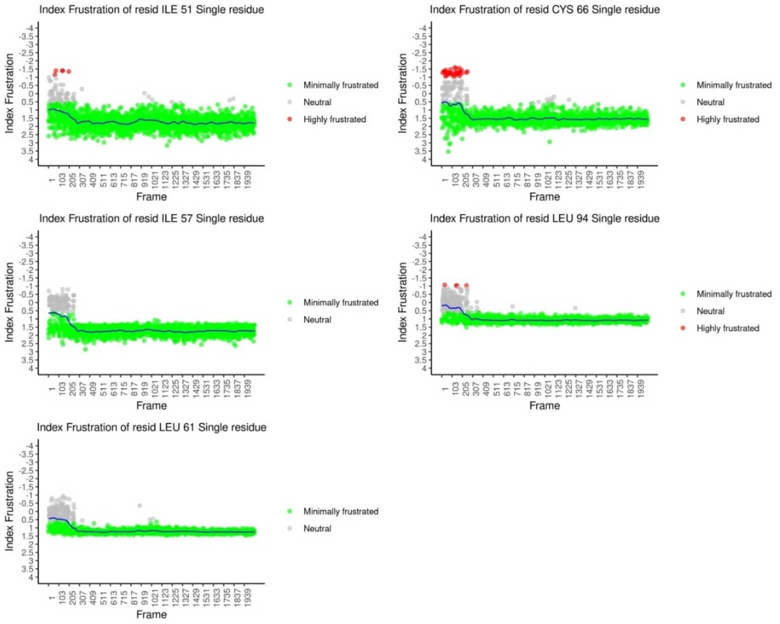


#### Cluster 4

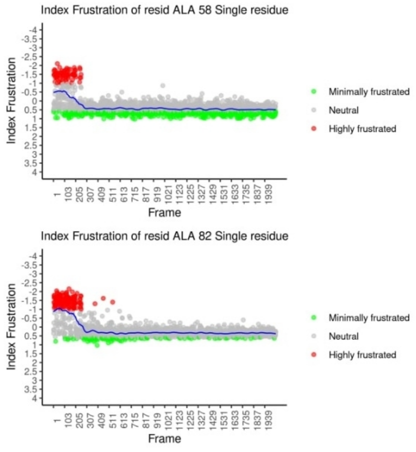

Shortly after the intermediate state is formed there exists an Expanded State 2 (E2) that gets stabilised thanks to further interactions with NfKB and internal interactions between Repeats 3 and 4. Residues in Cluster 5 are involved in these interactions.

#### Cluster 5

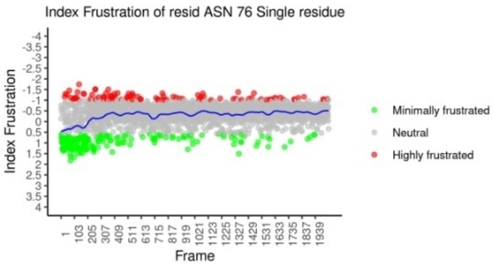


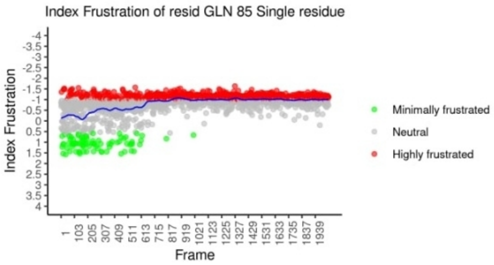


**2nd Folding Transition**

Clusters 3, 6 and 7 are similar to the ones previously described but they change their frustration state at a different point in time. These clusters have a transition from one frustration state to another approximately between frames 850 and 900 which corresponds to the region around Qw=0.45. This region comprises the barrier between the Expanded State2 (E2) and the Expanded State1 (E1) that is close to the Folded State (difference between them is that the N And C-terminal loop and b-hairpin regions of IkBa are frying around during the dynamics and "expand" the structure in its Rg and Qw coordinates). Clusters 3 and 6 are located in the helices from Repeats 5 and 6 and are important for their internal stability. Residues in Cluster 7 participate in the interaction with NFkB.

#### Cluster 3

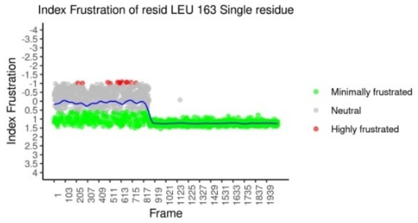


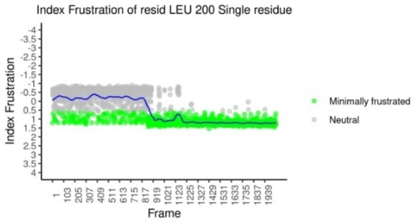


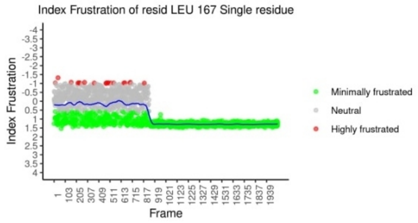


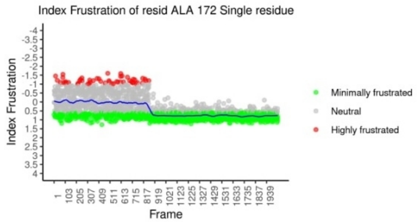


#### Cluster 6

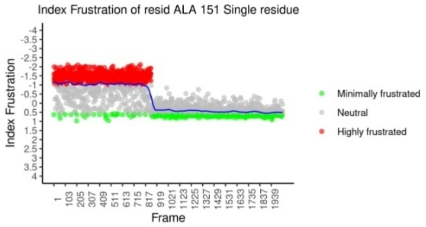


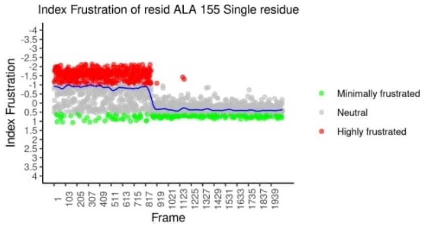

### Cluster 7

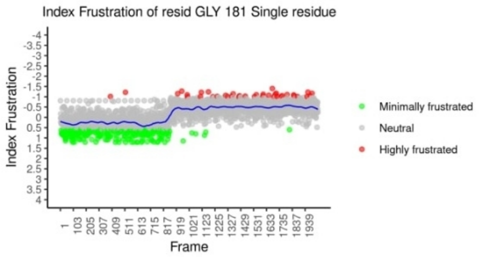


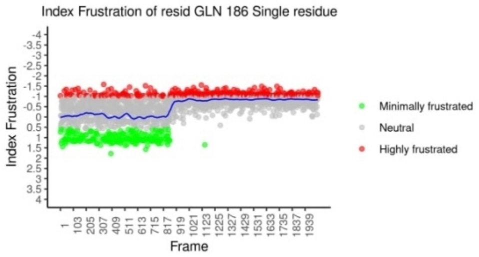


The previous panel graphics were obtained with the use of the function:

```
save_res_dynamic_clusters(Dynamic_sing, Clusters = "all", Nrow = 3, Ncol = 2)
```

The *Nrow* and *Ncol* parameters indicate the number of rows and columns per page of the stored PDFs.

#### Summarising

1. Calculate local frustration across dynamics with `dynamic_frustration()` using single-residue index (*Mode* = *"singleresidue"*).
2. Detect dynamic clusters with `detect_dynamic_clusters()` with the desired parameters.
3. View the results:
    - Filter in frustration mean and frustration dynamic range: `plot_variable_res_filter()`.
    - Fitted model of a specific residue: `plot_res_dynamics()`.
    - Undirected graph and clusters: `plot_dynamic_clusters_graph()`.
    - Pymol visualisation of clusters: `plot_clusters_pymol()`
    - Save `plot_res_dynamics()` graph of residuals in clusters: `save_res_dynamic_clusters()`.

#### Full script

#Load frustratometeR

```
library(frustratometeR)
```

### Defining the order of the structures

```
OrderList <- paste0("612_", 1:2040, ".pdb")
```

#Calculation of local energy frustration for all frames of dynamics

```
Dynamic_sing <- dynamic_frustration(PdbsDir = "/home/Dynamic/pdbs/", OrderList = OrderList, ResultsDir = "/home/Dynamic/results/", Mode = "singleresidue")
```

#Detect dynamic clusters

```
Dynamic_sing <- detect_dynamic_clusters(Dynamic = Dynamic_sing)
```

**Note: if your R language interpreter proposes the installation of Miniconda, we do not recommend its use in this case. Ideally use the version of python3 installed on your computer.**

#Visualisations

```
plot_variable_res_filter(Dynamic = Dynamic_sing)
```

```
plot_dynamic_clusters_graph(Dynamic = Dynamic_sing)
```

```
dynamic_res(Dynamic = Dynamic_sing, Resno = 85, Chain = "A")
```

```
dynamic_res(Dynamic = Dynamic_sing, Resno = 167, Chain = "A")
```

```
plot_res_dynamics(Dynamic = Dynamic_sing, Resno = 167, Chain = "A")
```

```
plot_res_dynamics(Dynamic = Dynamic_sing, Resno = 85, Chain = "A")
```

```
plot_clusters_pymol(Dynamic_sing, Clusters = "all")
```

```
save_res_dynamic_clusters(Dynamic_sing, Clusters = "all", Nrow = 3, Ncol = 2)
```

### 7. Running frustrations in a terminal or cluster

#### This example shows how to run frustratometeR from a terminal script

You can create a script which first loads frustratomeR and receives parameters through your OS terminal like this:

```
library(frustratometeR)
```

```
args = commandArgs(trailingOnly = TRUE)
```

The next step will depend on whether we want to create a script that calculates the local energy frustration of a protein structure per execution or multiple structures contained in the same directory. For the first case we will use the function `calculate_frustration()`, while for the second `dir_frustration()`.

##### For a protein structure ( `calculate_frustration()` )

Here the energy local frustration will be calculated for a structure contained in a file in PDB format ( *PdbFile*) in your computer or cluster.

#The received parameters are assigned to variables

```
PdbFile <- args[1]
```

```
Mode <- args[2]
```

```
Chain <- args[3]
```

```
Electrostatics_K <- args[4]
```

```
SeqDist <- args[5]
```

```
Graphics <- args[6]
```

```
Visualization <- args[7]
```

```
ResultsDir <- args[8]
```

#The data type of the Boolean variables is changed, since strings of characters are received by terminal

```
if(Chain == "NULL") Chain <- NULL
```

```
if(Electrostatics_K == "NULL") Electrostatics_K <- NULL
```

```
if(Graphics == "T") Graphics <- T else Graphics <- F
```

```
if(Visualization == "T") Visualization <- T else Visualization <- F
```

#Local energy frustration is calculated

```
calculate_frustration(PdbFile = PdbFile, Mode = Mode, Chain = Chain,  Electrostatics_K = Electrostatics_K, SeqDist = SeqDist,
ResultsDir = ResultsDir, Graphics = Graphics, Visualization = Visualization)
```

**Execution line example by terminal**

```
Rscript ourScript.R /home/1n0r.pdb configurational A NULL 12 T T /home/results/
```

##### For multiple protein structures ( `dir_frustration()` )

Here the energy local frustration is calculated for all the PDB format structure files contained in your computer or cluster that are in the same directory (*PdbsDir*).

#The received parameters are assigned to variables

```
PdbsDir <- args[1]
```

```
Mode <- args[2]
```

`Chain <- args[3]`

`Electrostatics_K <- args[4]`

`SeqDist <- args[5]`

`Graphics <- args[6]`

`Visualization <- args[7]`

`ResultsDir <- args[8]`

#The data type of the Boolean variables is changed, since strings of characters are received by terminal

`if(Chain == "NULL") Chain <- NULL`

`if(Electrostatics_K == "NULL") Electrostatics_K <- NULL`

`if(Graphics == "T") Graphics <- T else Graphics <- F`

`if(Visualization == "T") Visualization <- T else Visualization <- F`

#Local energy frustration is calculated

`dir_frustration(PdbsDir = PdbsDir, Mode = Mode, Chain = Chain,  Electrostatics_K = Electrostatics_K, SeqDist = SeqDist,  ResultsDir = ResultsDir, Graphics = Graphics, Visualization = Visualization)`

**Execution line example by terminal**

`Rscript ourScript.R /home/pdbs/ configurational A NULL 12 T T /home/results/`

#### Summarising

1. Load frustratometeR.
2. Receive parameters by console.
3. Control the data types of variables.
4. Use the necessary function, for example `calculate_frustration()` for one protein structure, `dir_frustration()` for multiple.
5. Execute the script with *Rscript* indicating the necessary parameters.

### 8. Docker container

For those users that for any reason are not able to install frustratometeR in their systems, we have generated a docker container so they can use the tool by command line. Be aware that this will only be able to calculate frustration values but will not be able to use the functions that display figures in RStudio or execute PyMol externally.

**1. Step: Install Docker:** https://docs.docker.com/

**2. Run your docker:** To calculate frustration for all PDBs that are contained in a specific folder in your computer, once docker is installed. You can download THIS BASH SCRIPT and execute it as a bash file in the following way:

sh frustra.sh **PDB_FOLDER FrustrationMode LicenceModeller**

Example: sh frustra.sh /Users/parra/Desktop/PDBs/test configurational XXXXXXXXXXXX

Since Modeller is an external tool that needs a licence to be used you have to provide its licence as the last parameter to run the script. Replace LicenceModeller with you valid Licence as the last parameter to use the docker script.

- You also have to change **FrustrationMode** by one of the 3 frustration indexes: configurational, mutational or singleresidue.
- Also you have to provide the Modeller Licence and replace **LicenceModeller** with the licence you get from Modeller.

### 9. Understanding frustration results

#### In `calculate_frustration()`

When frustratometeR calculates frustration a results folder is created (*ResultsDir*/jobID.done) for every structure (jobID). Users will find 3 subfolders with the results.

1) FrustrationData.
2) VisualizationScripts.
3) Images.
Here we describe all the files that are produced in each folder.

1) FrustrationData: This folder contains all the data that is calculated by frustratometeR. Depending on the frustration indexes that were selected to be calculated (i.e, configurational, mutational and singleresidue) you will find the following files:

```
*JobID.pdb_configurational: This file contains the "configurational frustration index" report for the contacts in the protein. One l
ine per contact. The columns in the file have the following meaning:

    Res1: Residue 1 in the interacting pair.
    Res2: Residue 2 in the interacting pair.
    ChainRes1: Chain to which the Residue 1 belongs in the PDB file (useful for multichain jobs).
    ChainRes2: Chain to which the Residue 2 belongs in the PDB file (useful for multichain jobs).
    DensityRes1: Local density for Residue 1.
    DensityRes2: Local density for Residue 2.
    AA1: Amino acid type for Residue 1.
    AA2: Amino acid type for Residue 2.
    NativeEnergy: Native Energy for the original interacting pair.
    DecoyEnergy: Mean value for the energy distribution calculated from the decoys.
    SDEnergy: Standard deviation for the energy distribution calculated from the decoys.
    FrstIndex: Configurational frustration index calculated for the interacting pair.
    Welltype: Type of contact based on the distance between the two residues and their local densities (long, short and water-mediat
ed).
    FrstState: Frustration class for the interacting pair.

*JobID.pdb_mutational: This file contains the "mutational frustration index" report for the contacts in the protein. One line per co
ntact. The file is structured in the same way as the JobID.pdb_configurational having the mutational frustration index value in the
FrstIndex column.

*JobID.pdb_singleresidue: This file contains the "single residue level frustration index" report. One line per residue. The file is
structured as follows:

    Res: Residue number.
    ChainRes: Chain to which the residue belongs.
    DensityRes: local density for the residue.
    AA: Amino acid type for the residue.
    NativeEnergy: Native energy for the residue.
    DecoyEnergy: Mean value for the energy distribution calculated from the decoys.
    SDEnergy: Standard deviation value for the energy distribution calculated from the decoys.
    FrstIndex: Single residue frustration index.

*JobID.pdb_configurational_5adens: This file contains the density of contacts around a sphere of 5 Armstrongs, centered in the C-alf
a atom from the residue. The different classes of contacts based on the configurational frustration index are counted both in absolu
te and relative terms. The file is structured as follows:

    Res: Residue number.
    ChainRes: Chain to which the residue belongs.
    Total: Total number of contacts within 5 Armstrongs from the C-alfa atom in the residue.
    nHighlyFrst: Fraction of highly frustrated contacts within 5 Armstrongs from the C-alfa atom in the residue.
    nNeutrallyFrst: Fraction of neutral contacts within 5 Armstrongs from the C-alfa atom in the residue.
    nMinimallyFrst: Fraction of minimally frustrated contacts within 5 Armstrongs from the C-alfa atom in the residue.
    relHighlyFrustrated: nHighlyFrst normalized to the Total Density.
    relNeutralFrustrated: nNeutrallyFrst normalized to the Total Density.
    relMinimallyFrustrated: nMinimallyFrst normalized to the Total Density.

*JobID.pdb_mutational_5adens: This file contains the density of contacts around a sphere of 5 Armstrongs, centered in the C-alfa ato
m from the residue. The different classes of contacts based on the mutational frustration index are counted both in absolute and re
lative terms. The file is structured in the same way as the *JobID.pdb_configurational_5adens file.
```

2) VisualizationScripts: This folder contains pymol and vmd scripts to visualize the frustration patterns over the protein structures. The following files are present in the folder.

```
*.pdb_configurational.tcl : a script to draw the 'configurational frustration
index' of contacts in the protein structure using VMD.

*.pdb_mutational.tcl:  a script to draw the 'mutational frustration
index' of contacts in the protein structure using VMD.

    To see the 3D structure with the contacts drawn you'll need a program
    called VMD (download free at http://www.ks.uiuc.edu/Research/vmd/ ).
    - open VMD, a command window will also open.
    - load the protein (go to File/NewMolecule and look for it in the
    folder were the Frustratometer results are)
    - then go to the command window and change directory to where the
    associated files for the pdb are, for example type in:
    "cd route_to_the_results_folder/frustratometer_jobID.pdb"
    - then type: source mfrst_yourprotein.pdb.tcl (this will actually draw
    the 'mutational frustration index' according to: highly frustrated
    contacts in red, the minimally frustrated in green, the direct ones in
    solid, and the water-mediated ones in dashed. "neutral" contacts are
    not drawn. All contacts are represented as lines emerging from the
    Calpha of each aminoacid)

    - you can also type from a console "vmd file.pdb -e file.tcl" replacing "file" with the name given to you jobid and then PyMOL will open and show the associated contacts.

*.pdb_configurational.pml : a script to draw the 'configurational frustration
index' of contacts in the protein structure using PyMOL.

*.pdb_mutational.pml:  a script to draw the 'mutational frustration
index' of contacts in the protein structure using PyMOL.

    To see the 3D structure with the contacts drawn using pml scripts you need a program called PyMOL.
    - open PyMOL, a command window will also open.
    - load the protein (go to File/Open and look for it in the folder where the FrustratometeR results are)
    - then go to File/Run.. and select the .pml file you wanna run.

    - you can also type from a console "pymol file.pml" replacing "file" with the name given to you jobid and then PyMOL will open and show the associated contacts.

*draw_links.py: This is a script needed to draw the frustratographs in PyMOL.
```

3) Images: This folder contains all the images that are generated by FrustratometeR when running `calculate_frustration()`.

#### In `mutate_res()`

As for the results of `mutate_res()`, these are stored in *ResultsDir*/jobId.done/MutationsData under the name of **FrustrationMode**_Res**Resno**_**MutationMethod**_**Chain**.txt.

In the case that the *Mode* is configurational or mutational, the file will contain the following columns corresponding to the frustration calculation for each of the 20 amino acid variants:

```
    Res1: Residue 1 in the interacting pair.
    Res2: Residue 2 in the interacting pair.
    ChainRes1: Chain to which the Residue 1 belongs in the PDB file (useful for multichain jobs).
    ChainRes2: Chain to which the Residue 2 belongs in the PDB file (useful for multichain jobs).
    AA1: Amino acid type for Residue 1.
    AA2: Amino acid type for Residue 2.
    FrstIndex: Configurational frustration index calculated for the interacting pair.
    FrstState: Frustration class for the interacting pair.
```

Instead, for single residue index:

```
    Res: Residue number.
    ChainRes: Chain to which the residue belongs.
    AA: Amino acid type for the residue.
    FrstIndex: Single residue frustration index.
```

On the other hand, saved graphics are stored in *ResultsDir*/jobId.done/MutationsData/Images

#### In `dynamic_frustration()`

The results of the `dynamic_frustration()` function are stored in the *ResultsDir/* directory where each folder corresponds to the result of `calculate_frustration()` for each frame. A file called Modes.log is stored in *ResultsDir*, which contains the frustration index used for the calculation. In this way, frustratometeR performs a check avoiding repeating the calculation, if necessary it can remove it.

By executing `dynamic_res()`, folders with the name Dynamic_plots_res_**Resno_Chain** are created in *ResultsDir*, which contains a file called **FrustrationMode**_Res_**Resno_Chain** which in each row has the data provided by `calculate_frustration()` in a certain frame for the *Resno* residue of the *Chain* chain. In addition, the corresponding graphics will be stored there.

**FrustratometeR in its execution clearly shows where the results were stored**
